# Inflammatory signals are sufficient to elicit TOX expression in mouse and human CD8 T cells

**DOI:** 10.1101/2021.03.15.435527

**Authors:** Nicholas J Maurice, Jacqueline Berner, Alexis K Taber, Dietmar Zehn, Martin Prlic

## Abstract

T cell receptor (TCR) stimulation leads to expression of the transcription factor TOX. Prolonged TCR signaling, such as encountered during chronic infections or in tumors, leads to sustained TOX expression, which induces a state of exhaustion or dysfunction. While CD8 memory T cells (T_mem_) in specific pathogen-free laboratory mice typically do not express TOX, functional human T_mem_ show heterogeneous TOX expression levels. Whether TCR-independent mechanisms can alter TOX expression in human and murine T_mem_ has not been defined. We report that human and mouse T_mem_ increase TOX expression following stimulation with inflammatory cytokines IL-12, IL-15, and IL-18. TOX and PD-1 expression patterns often appear to be directly correlated, however, we found that TOX is not necessary for cytokine-driven expression of PD-1. Together, these observations highlight that inflammation is sufficient to alter TOX and PD-1 expression and that the signals regulating TOX expression appear well conserved in human and murine T_mem_.

## Introduction

T cell exhaustion (i.e. dysfunction) is driven by chronic TCR stimulation with cognate antigen (Ag)^1, 2, 3^. It describes a differentiation state in which T cells have diminished capacity to respond to stimulatory inputs and limited effector capacity^2, 3, 4^. The purpose of T cell exhaustion during chronic infections may be to limit tissue pathologies when pathogen cannot be immunologically eliminated^5, 6^. Though exhaustion could be considered an immunologic concession during chronic infection, it also occurs in tumors and causes an attenuated anti-tumor cytotoxic T cell response^7^. Thus, mechanistically understanding and therapeutically overcoming T cell exhaustion has been a major goal of tumor immunotherapy. Chronic TCR stimulation elicits a program that leads to constitutively high expression of programmed cell death 1 (PD-1)^8^. PD-1 is an inhibitory receptor which is expressed by activated and exhausted T cells and often used as a biomarker to infer T cell functionality^9^. When bound to its ligands, PD-1 negatively regulates T cell function^2^. Therapeutic targeting of PD-1 with monoclonal antibodies, also referred to as immune checkpoint inhibitors, can reinvigorate a subset of these PD-1 expressing T cells^2,10,11,12^.

A set of recent studies demonstrated that the transcription factor, thymocyte selection-associated high mobility group box (TOX) protein, drives or stabilizes this TCR-mediated T cell dysfunction and PD-1 upregulation^6, 13, 14, 15, 16^. When stably expressed, TOX drove Ag-specific T cell exhaustion in mouse models of chronic lymphocytic choriomeningitis virus (LCMV) infection, transplantable B16 melanoma, and inducible hepatocellular carcinoma^6,13,14^. Further, putative tumor Ag-specific CD8 T cells isolated from primary human breast, ovarian, and skin cancer samples, as well as those specific for hepatitis C virus (HCV), mirrored this phenotype, suggesting TOX dictates exhaustion programs in humans, too^6, 13, 14^. Of note, TOX expression by HCV-specific T cells was reduced following treatment and clearance of the infection but still detectable at higher levels than in T cells from HCV infections that spontaneously resolved and among T cells specific for influenza A virus (IAV)^6^. Mechanistic insight was provided by targeted deletion of TOX in Ag-specific cytotoxic T cells, which diminished PD-1 expression and restored functionality at the expense of cell survival^6, 13^. Therefore, TOX concedes activation and effector function for exhaustion (i.e. PD-1 expression) and T cell survival during chronic TCR stimulation. In instances of brief TCR engagement, TOX is transiently induced to a level lower than that of exhausted T cells, but with limited known functional consequence^6, 13, 14^.

While the requirement for TOX has been well defined in the context of TCR-mediated dysfunction, there is nascent evidence that TOX expression by itself is not indicative of T cell exhaustion. Recent studies illustrated that TOX expression is detected in some functional CD8 memory T cells (T_mem_), for instance in CD8 effector memory (T_EM_) and effector memory CD45RA-expressing (T_EMRA_) subsets^17^. CD8 T_mem_ specific for the latent viruses, cytomegalovirus (CMV) and Epstein-Barr virus (EBV), had elevated TOX expression, compared to those specific for acute infections, which further suggests that TCR signals are critical in regulating TOX expression^17^. In a second study, it was shown that a fraction of the human T_mem_ population expresses *TOX* transcripts amongst other signature genes typically associated with T cell exhaustion^18^. The observation that functional humam memory T cells express TOX also led to questioning whether TOX is functionally conserved between mouse and human T cells^19^. Further complicating TOX and exhaustion, the murine tissue resident memory T cell (T_RM_) transcriptome is characterized by concomitant expression of transcripts encoding *Tox,* exhaustion markers, TCR signaling components, and cytotoxic molecules, well after initial priming events^20, 21^. While the role of TOX in these TOX-expressing populations with and without signs of T cell exhaustion is not fully understood, these data suggest that TOX expression by memory T cells cannot be reliably used to extrapolate T cell function.

While the role of TCR signals in initiating and maintaining PD-1 and TOX expression has been well established, relatively little remains knows about non-TCR signals that could regulate their expression in T cells^22^. We considered that cytokine-mediated stimuli could also affect TOX expression levels without promoting the induction of T cell exhaustion. First, pro-inflammatory cytokines, like IL-15, can induce PD-1 without agonist TCR signals. Second, T_RM_ that are likely not detecting cognate Ag still upregulate PD-1 and other markers associated with exhaustion^20, 21, 23, 24^, yet rely on IL-15 signaling for maintenance in some tissues^25, 26^. Thus, inflammatory signals could provide an explanation for some of the seemingly disparate results of TOX expression and T cell function. Here, we show that pro-inflammatory cytokines were sufficient to induce TOX expression in the absence of agonist TCR signals in both mouse and human CD8 T_mem_, while concurrently inducing expression of cytotoxic molecules. Together, these data demonstrate that TOX expression per se does not indicate TCR-mediated dysfunction or even a recent TCR signals. We also demonstrate that PD-1 expression is still upregulated in TOX-deficient T cells indicating that TOX is not necessary for PD-1 expression. Overall, our data reveal new TCR-independent mechanisms that shape TOX and PD-1 expression heterogeneity in T_mem_ and indicate that these mechanisms are conserved in both mouse and human T cells. Though these findings ultimately complicate the use of TOX exclusively as an exhaustion biomarker, they implicate TOX in inflammation-driven programs of memory T cell activation.

## Results

### Cytokine stimulation induces TOX expression in murine CD8 T_mem_

The pro-inflammatory cytokines IL-12, IL-15, and IL-18 elicit interferon-γ (IFNγ) and granzyme B (GzmB) expression in mouse and human CD8 T_mem_ in the absence of agonist TCR signals^27, 28, 29^. We first sought to determine if these cytokines could also induce TOX expression in a TCR-independent manner. To generate a well-defined population of CD8 T_mem_, we transferred congenically-marked OT-I CD8 T cells, which express a TCR specific for the SIINFEKL peptide of OVA, into wildtype C57BL/6J animals followed by infection with OVA-expressing vesicular stomatitis virus (VSV-OVA) (**Fig. 1a**). We waited ≥60 days before using these mice for subsequent experiments (referred to as VSV-OVA OT-I memory mice) **(Fig. 1a)**. We isolated T cells from the spleens and LNs from VSV-OVA OT-I memory mice using negative-selection magnet-activated cell sorting (MACS) prior to ex vivo stimulation experiments **(Fig. 1a)**. This was done to ensure that cytokines act directly on T cells^30^. As a negative control, we cultured bulk T cells in media alone (mock) and as a positive control, we stimulated T cells with anti-CD3/CD28 microbeads **(Fig. 1a)**. We used a combination of rIL-12, rIL-15, and rIL-18 (IL-12/15/18) to induce IFNγ and GzmB expression in a TCR-independent manner **(Fig. 1a).** We found that IL-12/15/18 stimulation induced PD-1 expression in OT-I T_mem_, but the increase in expression was markedly higher after TCR ligation **(Fig. 1b)**. PD-1 frequency and median fluorescence intensity (MedFI) in OT-I T_mem_ increased throughout the duration of IL-12/15/18 stimulation **(Fig. 1b)**. Similarly, TCR and IL-12/15/18 stimulation induced TOX upregulation in OT-I T_mem_ **(Fig. 1c)**. Next, we measured TCF1 expression, a transcription factor needed for memory T cell self renewal that is lost in terminally exhausted T_mem_^31, 32, 33, 34^. Alongside increasing PD-1 and TOX levels, both TCR- and IL-12/15/18-mediated stimulation led to significant loss of TCF1 expression in OT-I T_mem_ (**Fig. 1d**). In sum, these data indicate that phenotypes often associated with exhaustion can be induced by TCR-independent, cytokine-mediated T_mem_ activation. Finally, we sought to determine whether stimulation similarly affected endogenous CD8 T_mem_ and CD8 T_naïve_. IL-12/15/18 stimulation significantly increased TOX expression in endogenous CD8 T_mem_ but was not observed to the same degree in CD8 T_naïve_ (**Supplemental Fig. 1a, b**). This CD8 T_mem_-specific response is, too, reflected in IL-12/15/18-mediated upregulation of PD-1 (**Supplemental Fig. 1c, d**). This is likely, in some degree, to the different propensities of T cell subsets (both major and memory) to become efficiently activated by cytokines^35^ and differences in cytokine receptor expression (particularly naïve T cells, which require TCR-mediated activation to induce IL-12R and strongly increase IL-18R expression^36, 37^). Much akin to OT-I T_mem_, TCR stimulation dramatically increased both TOX MedFI and PD-1 expression across endogenous subsets (**Supplemental fig. 1a-d**), though the fold change in TOX staining intensity was most pronounced in CD8 T_mem_ (**Supplemental fig. 1a**). Though IL-12/15/18 stimulation increases TOX MedFI in transgenic and endogenous CD8 T_mem_, it is initially to a lower degree than that of TCR-stimulated cells (**Fig. 1c, Supplemental fig. 1a**). Since shortterm TCR- and IL-12/15/18-stimulation could dramatically augment TOX and PD-1 expression in CD8 T_mem_ from VSV-OVA OT-I memory mice, we next sought to test if TOX and PD-1 upregulation compromises functionality.

**Figure 1.**
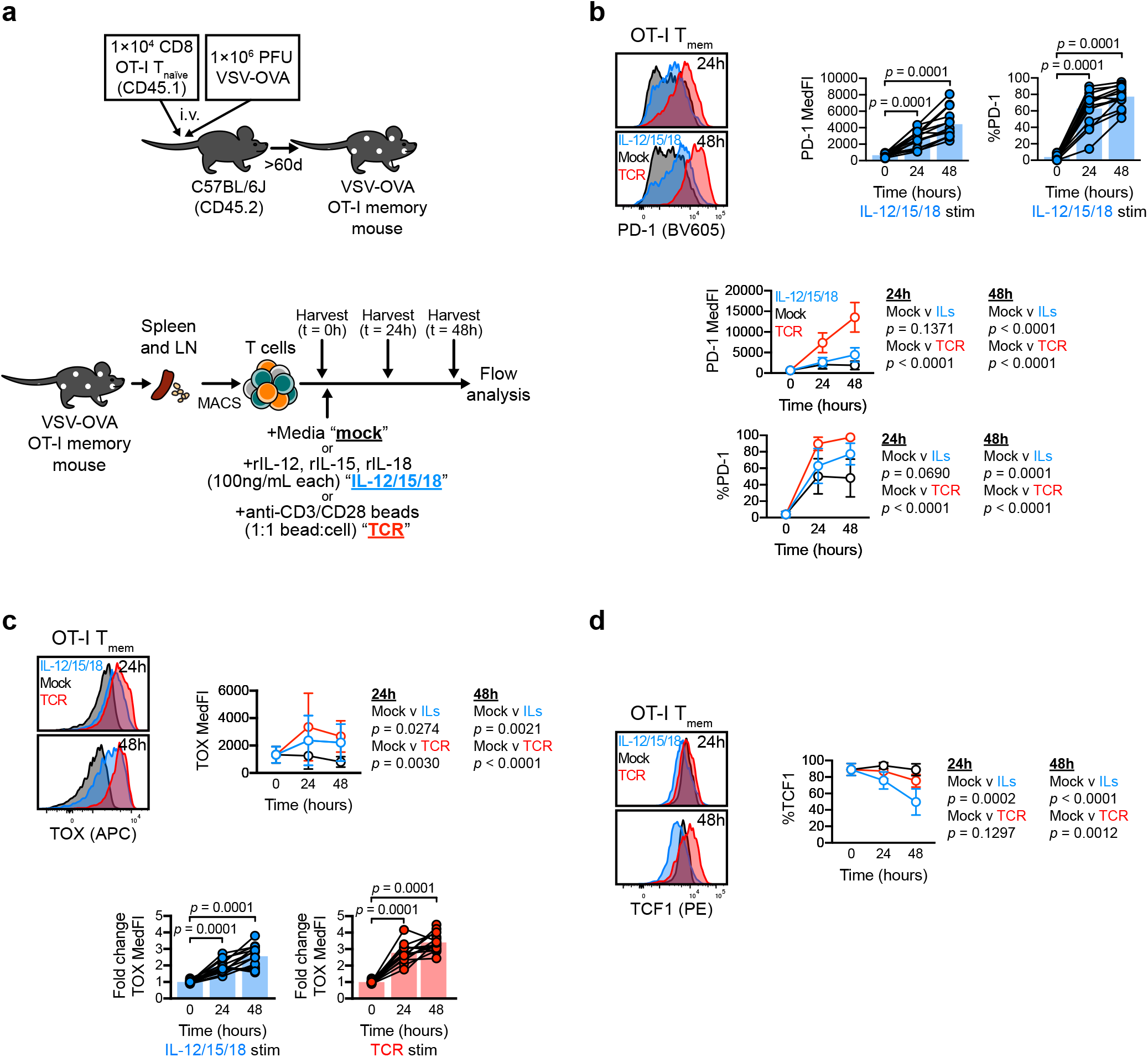
Cytokine stimulation induces TOX expression in murine CD8 T_mem_. **a** Schematic of OT-I memory mouse generation (top) and subsequent stimulation assays (bottom). OT-I T_naïve_ were transfered and expanded with VSV-OVA, then aged to stable memory contraction; after, T cells were enriched from VSV-OVA expanded OT-I memory animals and stimulated with media alone (mock), IL-12, −15, and −18 in combination (IL-12/15/18) (each at 100ng/mL), or anti-CD3/CD28 microbeads (TCR) at a ~1:1 bead:cell ratio. **b-c** expression of **b** PD-1, **c** TOX, and **d** TCF1 within stimulated OT-I T_mem_ throughout experiment time course. TOX MedFI fold change in **c** was calculated against average TOX MedFI from mock stimulations in a subset-specific, batch-specific, and timepointspecific manner. In **b** and **c**, bar chart symbols represent one animal at a unique timepoint/condition and are connected by animal identity, with bar indicating mean; the indicated statistical significances were calculated using Wilcoxon matched-pairs signed rank tests. In **b**-**d**, symbols in line plots comparing stimulation conditions represent the mean across all animals for a specific timepoint/condition ± SD; the indicated statistical significances were calculated using Mann-Whitney tests. Figures in **b** and **c** depict results from *n* = 14 mice across 7 experiments. Figures in **d** depict results from *n* = 9 mice across 2 experiments. All representative flow plots are sourced from the same animal.

### Functional CD8 T_mem_ express TOX and PD-1 and effector proteins

We isolated T cells from VSV-OVA OT-I memory mice as outlined for Fig. 1. We stimulated T cells in the presence of Golgi inhibitors and found that OT-I T_mem_ produced substantial amounts of IFNy after IL-12/15/18 or TCR stimulation(**Fig. 2a**); yet IFNγ-expressing OT-I T_mem_ demonstrated higher TOX and PD-1 expression than those that failed to make IFNγ (**Fig. 2b, c**). Similarly, OT-I T_mem_ that produced GzmB post-stimulation also demonstrated increased TOX and PD-1 expression (**Supplemental fig. 2a, b**). Together, these data indicate that TOX and PD-1 expression are elevated in activated, functional CD8 T_mem_ and suggest that TOX expression is also part of a cytokine-driven T cell activation program.

**Figure 2.**
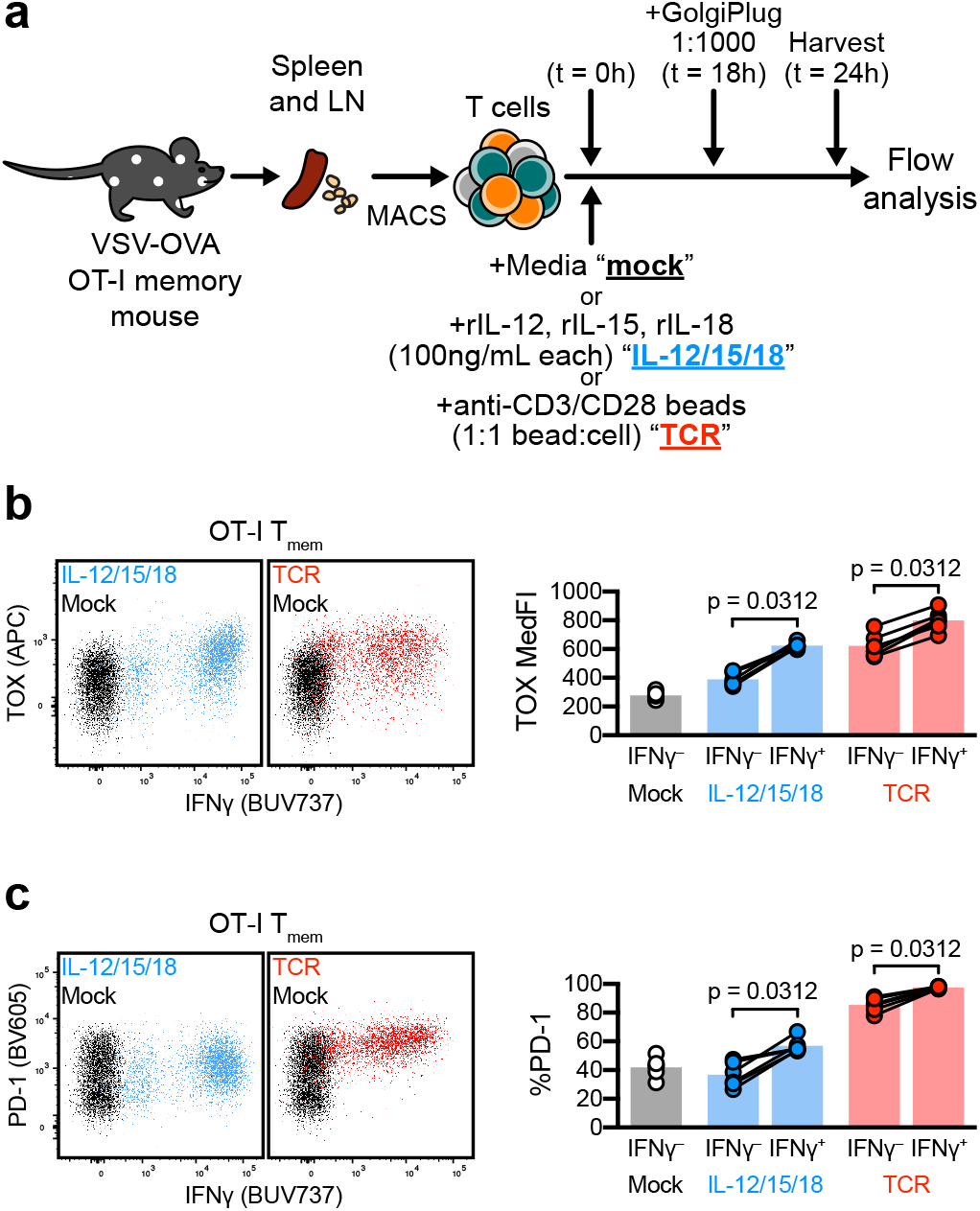
TOX and PD-1 expression occur in functional CD8 T cells. **a-c** Intracellular cytokine staining (ICS) in tandem with TOX interrogation. **a** Experiment schematic, in which bulk T cells from VSV-OVA OT-I memory mice were stimulated (mock, black; IL-12/15/18, blue; TCR, red). Cells were treated with GolgiPlug 18h into stimulation and harvested for flow staining and analysis at 24h. **b, c** Expression of **b** TOX and **c** PD-1 in IFNγ^+^ and IFNγ^-^ OT-I T_mem_. Representative plots depict cells from the same animal across different stimulation conditions. Symbols in **b** and **c** represent a T cell population within a unique animal with symbols connected by animal identity (*n* = 6 across 2 experiments). Bars represent mean and indicated statistical significances were calculated by Wilcoxon matched-pairs signed rank test.

### Induction of TOX and PD-1 is heterogeneous in CD8 T_mem_

To ensure that our data were not solely reliant on OT-I T cells, we also generated gBT-I memory mice using gBT-I TCR transgenic cells (specific for an epitope of the HSV2 gB protein) and a recombinant, gB epitope-expressing LM strain (LM-gB) (**Supplemental Fig. 3b**). After stable contraction of TCR transgenic T_mem_ (≥60d), we conducted stimulation assays as previously outlined (**Fig. 1a**). IL-12/15/18- or TCR-mediated stimulation led to comparable TOX upregulation in OT-I and gBT-I T_mem_ (**Fig 3a, b**). Similarly, PD-1 expression was comparable in OT-I and gBT-I T_mem_ after stimulation (**Supplemental Fig 3c, d**), with a concurrent loss of TCF1 expression (**Supplemental Fig. 3e, f**). We next asked if altering the nature of the priming infection could affect the ability to express TOX in response to cytokine-mediated activation at the memory stage. We adoptively transferred P14 transgenic T cells, a TCR transgenic specific for lymphocytic choriomeningitis virus (LCMV) gp33, followed by infection with LCMV Armstrong or Docile (**Supplemental Fig. 3g, h**). These LCMV strains elicit acute and chronic infections, respectively (the latter causing T cell dysfunction). We then stimulated (same culture set-up as outlined in **Fig. 1a**) T cells from these P14 memory mice. P14 T_mem_ from LCMV Armstrong infected mice readily upregulated PD-1 after TCR- or IL-12/15/18 stimulation (**Supplemental Fig. 3i**). The exhausted P14 T_mem_ from LCMV Docile-infected mice already uniformly expressed PD-1 prior to stimulation; but IL-12/15/18 or TCR stimulation further increased surface PD-1 expression (via increased MedFI) (**Supplemental Fig. 3j**). P14 T_mem_ from LCMV Armstrong infected mice increased TOX expression after TCR or IL-12/15/18 stimulation (**Fig. 3c**). However, exhausted P14 T_mem_ from LCMV Docile-infected mice only significantly increased TOX expression after TCR stimulation (**Fig. 3d**) and showed significantly lower fold changes in TOX MedFI compared to P14 T_mem_ from LCMV Armstrong-infected mice. While differences between CD8 T_mem_ from acute and chronic infected are expected, the differences between gBT-I and OT-I (~3 to 4-fold increase in TOX expression) compared to P14 (up to ~2-fold) need to be interpreted with caution since the gBT-I, OT-I and P14 experiments used different TOX antibody clones (REA473 and TXRX10, respectively). Overall, our data indicate that T_mem_ that were generated by different acute infections increase TOX expression in response to pro-inflammatory cytokines suggesting that this a broadly applicable mechanism of TOX induction in the memory T cell compartment. We next sought to determine if PD-1 and TOX upregulation in response to stimulation was similarly recapitulated in human CD8 T cells.

**Figure 3.**
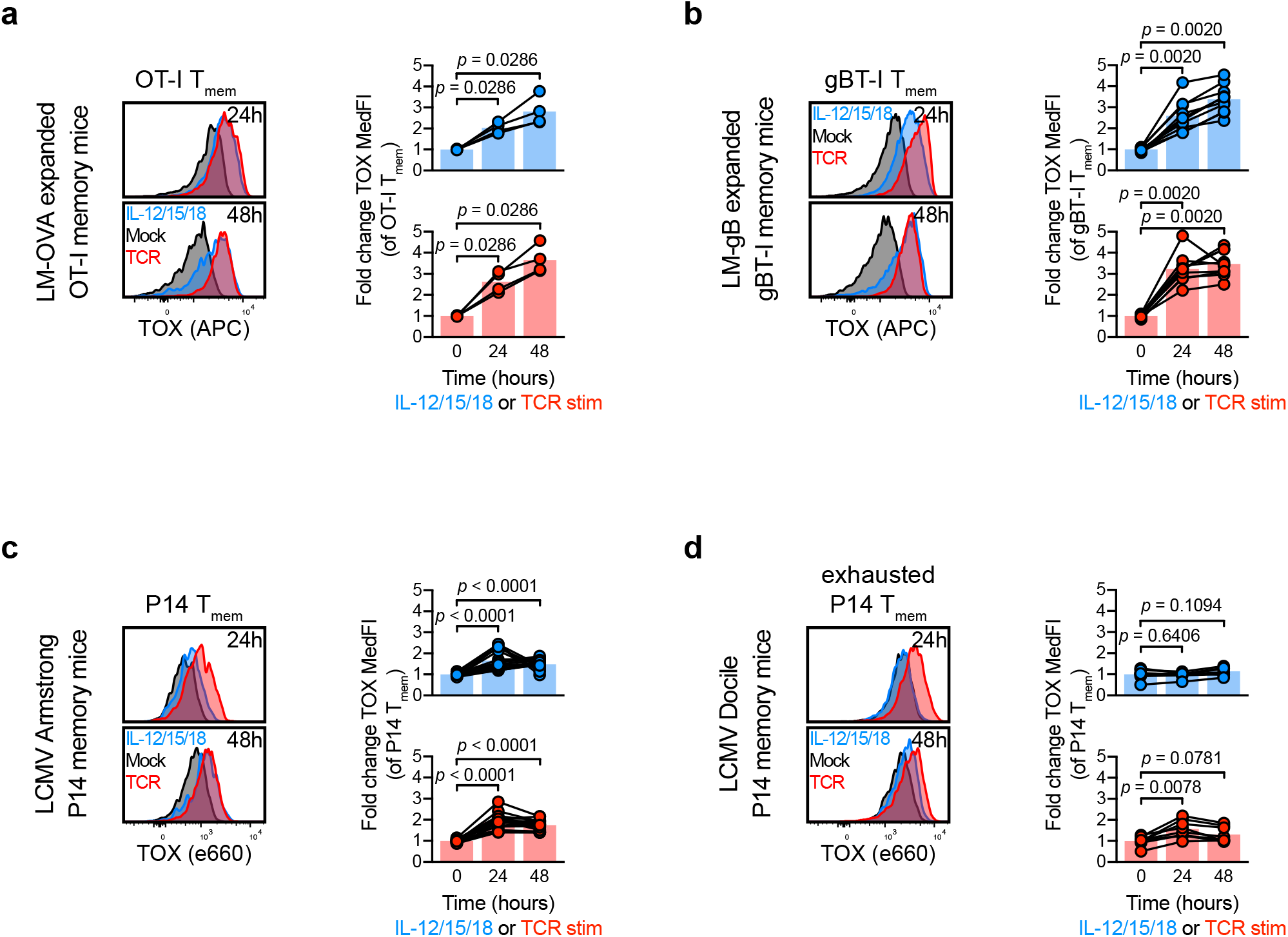
Cytokine-mediated TOX induction is limited in exhausted T cells. **a-b** Changes in TOX expression within LM-expanded TCR transgenic T_mem_: OT-I, specific for OVA Ag and gBT-I, specific for gB Ag. MACS-enriched T cells from LM-expanded OT-I or gBT-I memory mice were stimulated with media alone (mock), recombinant IL-12, −15, and −18 in combination (IL-12/15/18) (each at 100ng/mL), or anti-CD3/CD28 microbeads (TCR) at a ~1:1 cell:bead ratio. **a, b** Representative TOX expression and TOX MedFI fold change during stimulation in LM-primed **a** OT-I and **b** gBT-I T_mem_. **c-d** Changes in TOX expression within LCMV-specific TCR transgenic P14 T cells expanded by acute (Armstrong, Arm.) or chronic (Docile, Doc.) LCMV infection. **c, d** Representative TOX expression and TOX MedFI fold change during stimulation in P14 T cells primed by **c** LCMV Armstrong and **d** LCMV Docile. TOX MedFI fold change in **a-d** was calculated against average TOX MedFI within mock stimulation in a batch-specific, timepoint-specific manner. We calculated indicated statistical significances in **a-d** using Wilcoxon matched-pairs signed rank tests. Each symbol in **a-d** represents a sample at a unique timepoint/condition, with bars delineating mean, which are connected by donor in **a-d** (*n* = 4 LM-OVA expanded OT-I memory mice across 2 experiments; *n* = 10 LM-gB expanded gBT-I memory mice across 2 experiments; *n* = 17 LCMV Armstrong-expanded P14 memory mice across 4 experiments; *n* = 8 LCMV Docile-expanded P14 memory mice across 2 experiments). Mouse identities are consistent between representative flow plots within the same generation/adoptive transfer condition.

### Cytokine stimulation induces TOX and PD-1 in human T_mem_

Using cryopreserved PBMCs from healthy, HIV-seronegative donors, we interrogated TOX and PD-1 expression by flow cytometry. We specifically gated CD8 T cells by a memory and naïve binary, delineating CD8 T_naïve_ as CD45RO-negative CCR7-positive, with remaining cells as CD8 T_mem_^38^ (**Fig. 4a**), and interrogated basal TOX and PD-1 expression between these two subsets (**Fig. 4a**). Since PD-1 expression is heterogeneous in humans^39, 40^, we measured TOX MedFI across PD-1 low-, medium-, and high-expressing events. We found that CD8 T_mem_ with the highest PD-1 expression also demonstrated significantly elevated TOX MedFI (**Fig 4b**), mirroring correlations of TOX and PD-1 expression in our mouse model as well as human HCV infections^6^. We next tested whether IL-12/15/18 stimulation increases PD-1 and TOX expression in T cell subsets and included mock and TCR stimulation conditions as negative and positive controls, respectively. We also included stimulations using rIL-6, rIL-15, or rIL-12 and rIL-18. We chose these additional conditions as IL-6 activates CD8 T_naïve_ (as evidenced by CD69 upregulation) and to discern individual activating contributions of each cytokine (**Supplemental Fig. 4a**). Across these conditions, IL-12/15/18- and TCR-mediated stimulations led to the most prominent increase of TOX staining intensity and PD-1^hi^ frequency in CD8 T_mem_ (**Fig. 4d**). We measured TCF1 expression after mock, IL-12/15/18, and TCR stimulation. A decrease in TCF1 expression accompanied an increase in TOX and PD-1 expression after IL-12/15/18 or TCR stimulation (**Supplemental fig. 4b**), akin to our mouse stimulation data. We further tested the degree of similarity between human and mouse T cells by measuring PD-1, TCF1, and TOX expression profiles in stimulated human CD8 T_naïve_. Like mouse CD8 T_naïve_, only TCR stimulation could lead to appreciable changes in TOX and PD-1 within human CD8 T_naïve_ (**Fig. 4c, Supplemental fig. 4c**). Since IL-6 can activate CD8 T_naïve_, we used this condition to determine if PD-1 and TOX expression could occur in naïve T cells in the absence of a TCR signal. Despite inducing CD69 expression, we found that IL-6-mediated stimulation failed to increase TOX or PD-1 expression in CD8 T_naïve_ (**Supplemental fig. 4d**). Together, these data show that CD8 T_mem_ differentially expressed TOX, PD-1, and TCF1 at homeostasis and after both IL-12/15/18 and TCR stimulation. We next wanted to better define these changes across different memory T cell subsets.

**Figure 4.**
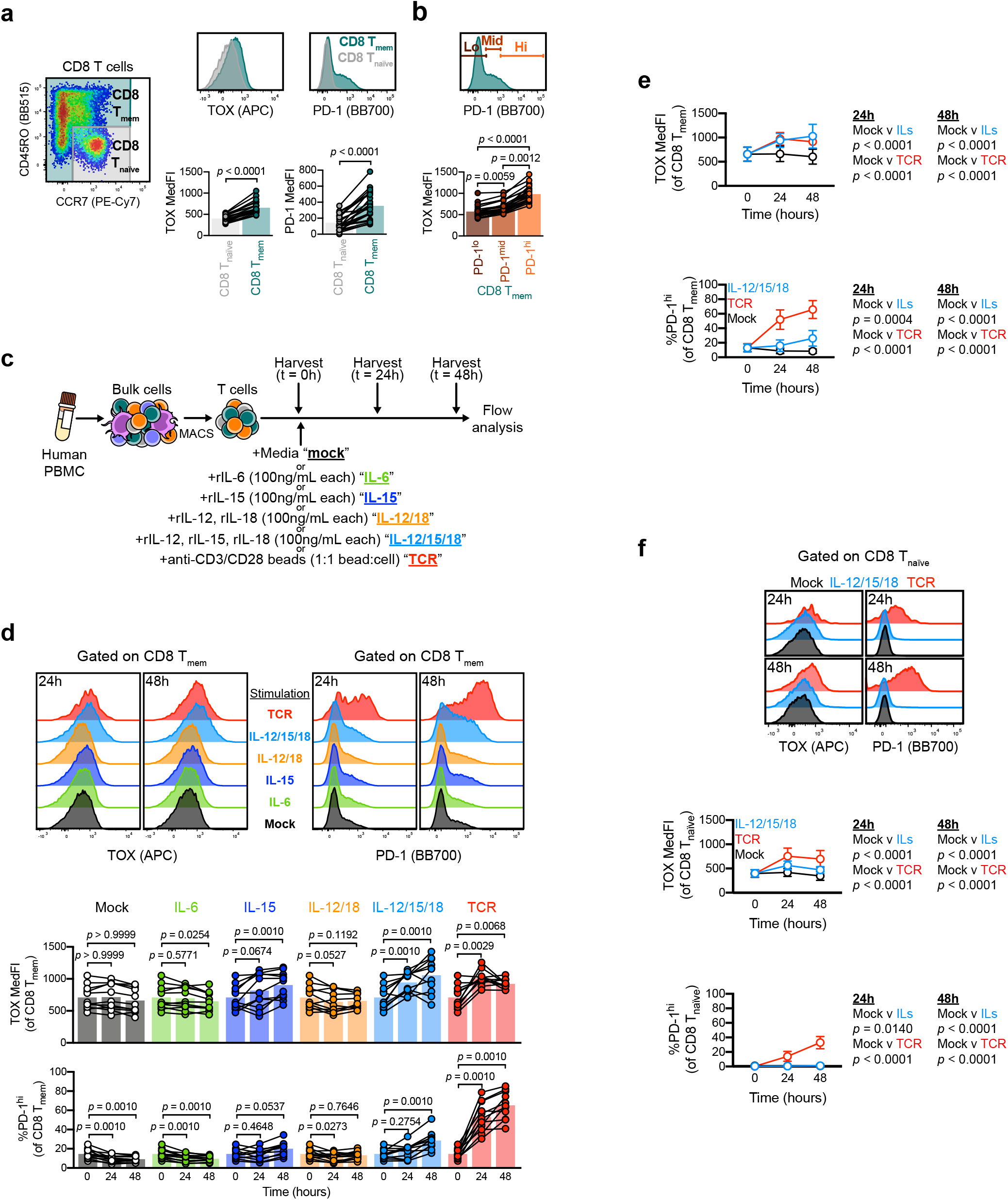
Inflammatory cytokines are potent inducers of TOX and PD-1 in human T_mem_. **a** Basal expression of TOX and PD-1 in CD8 T_mem_ and T_naïve_. **b** TOX MedFI across PD-1 low, medium, and high expressing CD8 T_mem_. **c** Schematic detailing T cell isolation from cryopreserved PBMCs and subsequent stimulation with recombinant IL-6, IL-15, IL-12 and −18, IL-12 and −15 and −18 (all at 100ng/mL, each), or anti-CD3/CD28 microbeads (TCR, 1:1 bead to cell ratio) and subsequent flow interrogation. **d** TOX expression (MedFI) and PD-1^hi^ frequency in CD8 T_mem_ throughout stimulation time course. **e, f** Comparison of TOX MedFI and PD-1^hi^ frequency in mock-, IL-12/15/18-, and TCR-stimulated **e** CD8 T_mem_ and **f** CD8 T_naïve_. In **a, b, d, e, f** we calculated indicated statistical significances by **a, d** Wilcoxon matched-pairs signed rank tests, **b** Friedman test with Dunn’s multiple comparisons tests, or **e, f** Mann-Whitney tests. In **a, d** each symbol represents a unique timepoint/treatment connected by donor with bars indicating mean **a** (*n* = 23 across four experiments) **d** (*n* = 11 across two experiments). In **e, f** each symbol represents the mean ± SD of the stimulation condition from *n* = 23 donors across four experiments. Representative plots from **a**, **d**, **f** are sourced from the same donor.

### Inflammation-induced PD-1 and TOX expression occur in most but not all CD8 T_mem_ subsets

To test if inflammation-induced PD-1 and TOX expression differs across human CD8 T_mem_ subsets, we used CD45RO and CCR7 staining to further delineate central memory (T_CM_) (CD45RO^+^ CCR7^+^), T_EM_ (CD45RO^+^ CCR7^-^), and T_EMRA_ (CD45RO^-^ CCR7^-^) subsets^38,41^ (**Fig. 5a**). When we measured TOX, PD-1, and TCF1 expression across these subsets, we noted that a substantial fraction of CD8 T_EM_ events were PD-1^hi^, and both CD8 T_EM_ and T_EMRA_ expressed elevated and lower levels of TOX and TCF1, respectively, at homeostasis (**Fig. 5b**). While this observation is in line with the initial report demonstrating TOX heterogeneity in human CD8 T_mem_ subsets^17^, it remained unknown if these CD8 T_mem_ subsets are equally capable of further TOX upregulation after stimulation. We observed that TOX, PD-1, and TCF1 expression kinetics in CD8 T_CM_ and T_EM_ largely resembled one another, with both IL-12/15/18 and TCR stimulation increasing the frequency of PD-1^hi^ events and TOX MedFI, but decreasing TCF1 MedFI (**Fig. 5c**). It is worth noting, that while TCF1 MedFI in CD8 T_CM_ drops profoundly after IL-12/15/18 or TCR stimulation, the loss in frequency of TCF1-expressing cells (as defined by subjective gating) is not as pronounced as what we observed in CD8 T_EM_ (**Supplemental fig. 5a**). While IL-12/15/18- and TCR-mediated stimulation were both able to significantly increase the frequency of PD-1^hi^ events and lower TCF1 MedFI in CD8 T_EMRA_, the degree of these changes was less pronounced than in CD8 T_CM_ or T_EM_ (**Fig. 5c**). Moreover, CD8 T_EMRA_ did not significantly upregulate TOX expression after TCR stimulation. This, however, was not due to inability to be stimulated, as CD8 T_EMRA_ readily expressed the activation marker CD69 after cytokine- or TCR-mediated stimulation (**Supplemental Fig. 5a**). Finally, it is worth noting that when stimulated with IL-15 alone, CD8 T_CM_, unlike CD8 T_EM_ and T_EMRA_, fail to significantly express PD-1 (**Supplemental fig. 5b**).

**Figure 5.**
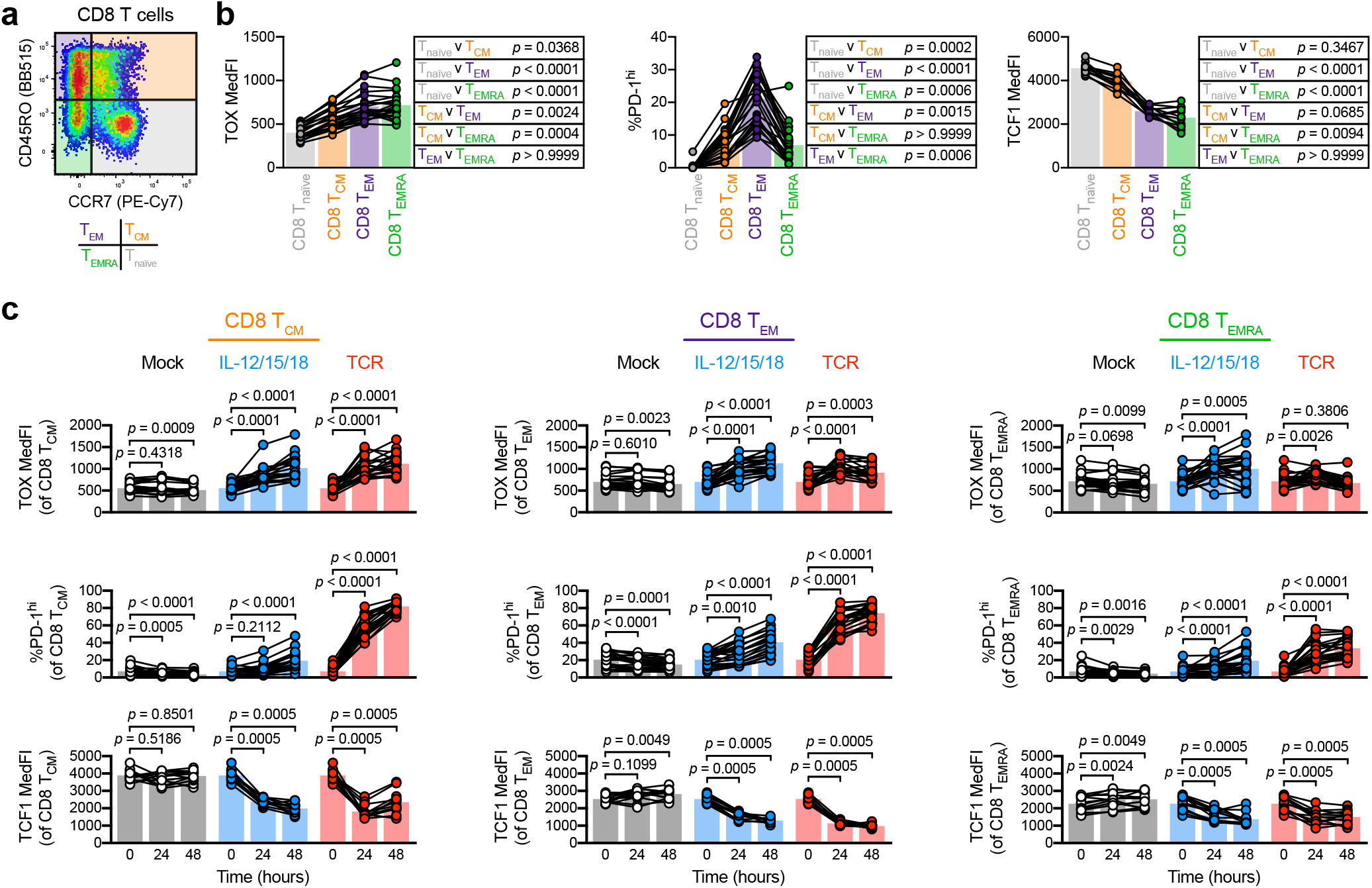
TOX and PD-1 upregulation are largely independent of T_mem_ subset. **a-c** Basal and stimulation-induced TOX and PD-1 expression in CD8 memory subsets. **a** Representative gating of CD8 T cells into naïve (T_naïve_, grey), central memory (T_CM_, orange), effector memory (T_EM_, purple), and effector memory CD45RA-expressing (T_EMRA_, green) subsets. **b** basal expression levels (MedFI) of TOX and TCF1 and frequency of PD-1^hi^ cells across CD8 T cell memory subsets. **c** TOX MedFI, PD-1^hi^ frequency, and TCF1 MedFI after mock (black), IL-12/15/18 (each at 100ng/mL, blue), or TCR (1:1 bead to cell ratio, red) stimulation in CD8 T_CM_ (left column), CD8 T_EM_ (center column), and CD8 T_EMRA_ (right column). Symbols in **b** and **c** represent unique samples (by timepoint/condition/subset) and are connected by donor identity, with bars representing mean. We determined statistical significances in **b** and **c** respectively using Friedman tests and Wilcoxon matched-pairs signed rank tests. **b** and **c** depict *n* = 23 donors across four experiments, except for TCF1 plots, which depict *n* = 12 donors across two experiments.

We next interrogated T cells with defined TCR specificity, specifically influenza A virus (IAV)-specific CD8 T cells using HLA-A*02 tetramers loaded with the GILGFVFTL peptide (**Fig. 6a**). We examined this CD8 T_mem_ population because these cells were reported to not express appreciable levels of TOX at homeostasis, likely owing to their T_CM_ phenotype^17^. Within our sample set, IAV-specific CD8 T cells were predominantly T_CM_ in half of the HLA-A*02 PBMC donors (**Fig. 6a**). Nevertheless, all IAV-specific CD8 T cells were able to substantially upregulate TOX and PD-1 expression after IL-12/15/18 stimulation (**Fig. 6b**), indicating that CD8 T_mem_ low for TOX and PD-1 at homeostasis, can also contribute to TOX and PD-1 heterogeneity after recent activation. Alongside testing IAV-specific CD8 T cells, we also interrogated the effects of stimulation in mucosal associated invariant T (MAIT) cells. We selected this population because 1) MAIT cells are non-conventional T cells, recognizing bacterial metabolites as Ags presented on MHC-related 1 (MR1)^42^, 2) inflammation is necessary for sustained MAIT cell effector function^43, 44^ and 3) MAIT cells are near-uniformly T_EM_ when defined by CD45RO and CCR7^45^. We identified MAIT cells using MR1 tetramers loaded with the 5-OP-RU metabolite^46^, which largely fell into our T_EM_ gate (**Fig. 6c**). Like IAV-specific CD8 T cells, IL-12/15/18 stimulation led to substantial TOX and PD-1 upregulation in MAIT cells (**Fig. 6d**). Since inflammation is necessary for sustained MAIT cell effector function, we asked if MAIT cells are differentially capable of responding to other cytokine combinations. Alongside IL-12/15/18, IL-15 alone, or IL-12 and IL-18 in unison could significantly increase both the PD-1^hi^ frequency and TOX MedFI of MAIT cells, but not IAV-specific T cells (**Supplemental fig. 6a, b**). Together, these data indicate that this cytokine-driven activation program is conserved across conventional and innate-like T cells.

**Figure 6.**
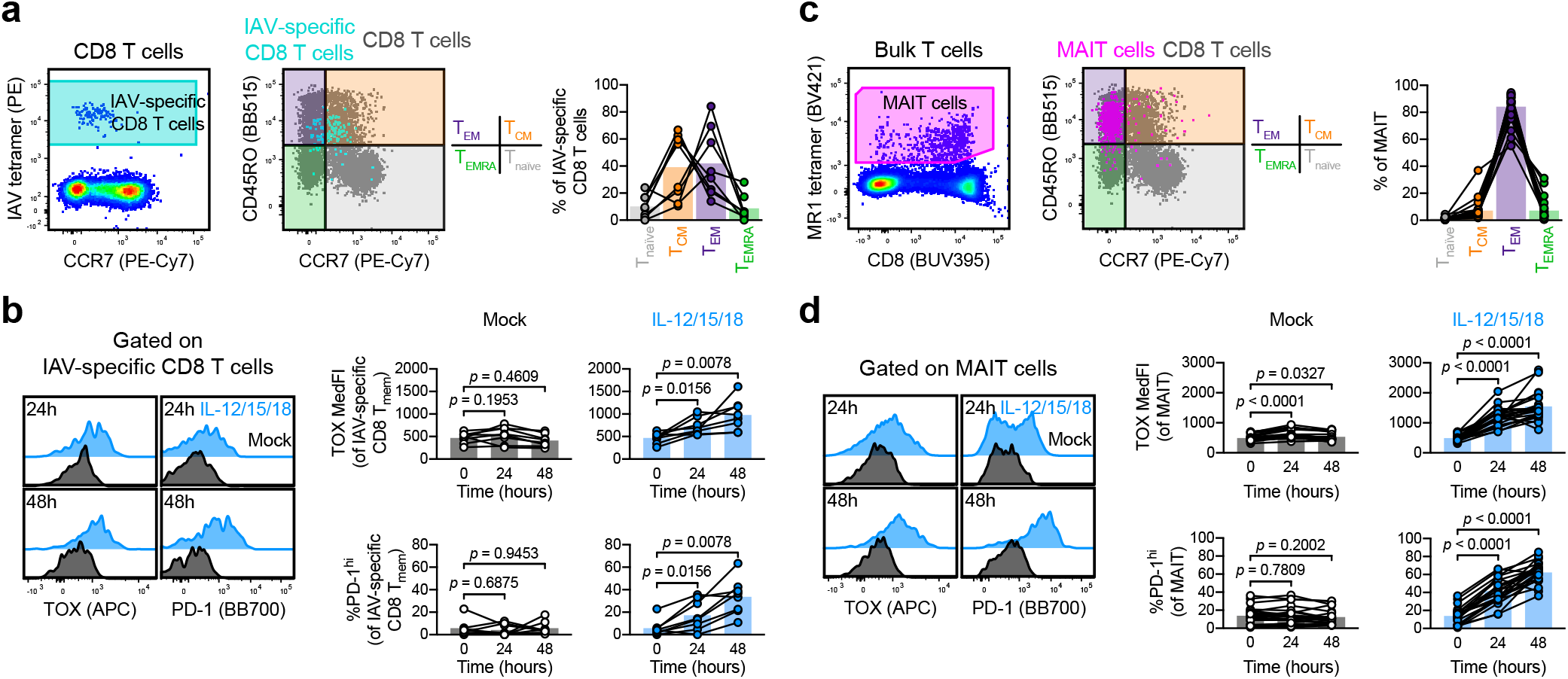
Stimulation induces TOX and PD-1 expression in conventional and innate-like T cells. **a-b** TOX and PD-1 induction in influenza A virus (IAV)-specific CD8 T cells. **a** Gating and memory phenotyping of IAV-specific CD8 T cells. **b** Induction of TOX and PD-1 in IAV-specific CD8 T cells by mock (black) or IL-12/15/18 (each at 100ng/mL, blue) stimulation. **c-d** TOX and PD-1 induction in mucosal associated invariant T (MAIT) cells. **c** Gating and memory phenotyping of MAIT cells. **d** Induction of TOX and PD-1 in MAIT cells by mock (black) or IL-12/15/18 (each at 100ng/mL, blue) stimulation. Representative plots in **a-d** are sourced from the same donor. Symbols in **a-d** represent unique samples (by timepoint/condition/subset) and are connected by donor identity, with bars representing mean. We determined statistical significances in **b** and **d** using Wilcoxon matched-pairs signed rank tests. **a** and **b** depict *n* = 8 donors across two experiments; **c** and **d** depict *n* = 23 donors across four experiments.

### Cytokine stimulation-induced PD-1 expression is independent of TOX

Finally, since PD-1 and TOX upregulation appeared tightly associated following cytokine-driven activation, we next asked if this association is mechanistic in nature. If TOX is necessary for PD-1 expression, it would allow to use the surface-expressed PD-1 as a surrogate for the intracellularly expressed TOX. TOX expression appears to drive PD-1 expression in a number of contexts, as exhausted T_mem_ dramatically downregulated PD-1 after TOX deletion or knockdown^6, 13, 15, 47^. Conversely, T cell transduction with TOX-encoding constructs leads to PD-1 upregulation^13, 14, 15, 47^. While TOX controls PD-1 expression during exhaustion, the role of TOX is less clear in activation. To dissect the function of TOX in stimulation-mediated PD-1 upregulation, we used wildtype (WT) and *Tox^-/-^* P14 T_mem_. To generate these P14 T_mem_, we adoptively transferred wildtype or knockout P14 T cells into C57BL/6J hosts, which we subsequently infected with LCMV Armstrong to form a T_mem_ population (**Supplemental fig. 7a**). To determine if TOX deficiency alters stimulation-induced PD-1 upregulation, we cultured MACS-isolated T cells from WT and *Tox^-/-^* P14 memory mice (28 days post LCMV Armstrong infection) in the presence of mock, IL-12/15/18, or TCR stimulation (**Supplemental fig. 7a**). Both WT and *Tox^-/-^* P14 T_mem_ increased PD-1 expression after IL-12/15/18 or TCR stimulation (**Fig. 7a, b, c**). Together these data indicate that TOX alone is not necessary for PD-1 upregulation in cytokine-stimulated CD8 T_mem_ and suggest other transcription factors are sufficient to drive PD-1 expression in the absence of TOX.

**Figure 7.**
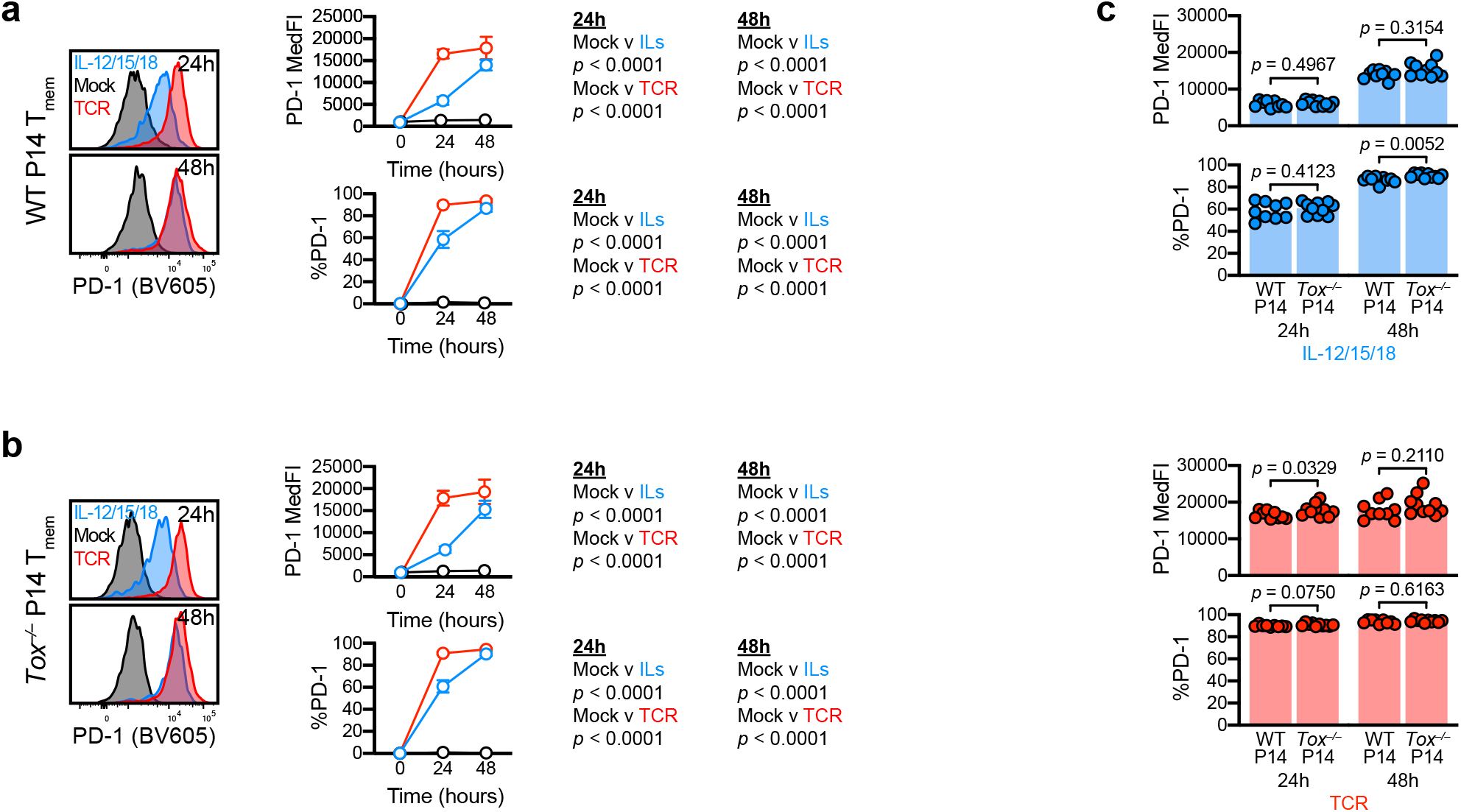
TOX deficiency does not abrogate stimulation-induced PD-1 expression. **a-c** Stimulation-induced PD-1 expression in WT and *Tox^-/-^* P14 T_mem_. T cells were stimulated with media alone (mock), recombinant IL-12, −15, and −18 in combination (IL-12/15/18 or ILs) (each at 100ng/mL), or with anti-CD3/CD28 microbeads at an ~1:1 cell:bead ratio (TCR). **a, b** PD-1 MedFI and expression frequencies in **a** WT or **b** *Tox^-/-^* P14 T_mem_ over stimulation time course. **c** Comparison of PD-1 MedFI and expression frequencies between IL-12/15/18 (left) or TCR (right) stimulated WT and *Tox^-/-^* P14 T_mem_. All indicated statistical significances were calculated using Mann-Whitney tests. Symbols in **a** and **b** represent the mean ± SD from all animals at a specific time/condition; and symbols in **c** represent stimulated P14 T_mem_ populations within a single animal (*n* = 9 WT P14 recipients and *n* = 10 *Tox^-/-^* P14 recipient across 2 experiments).

## Discussion

TOX has been foremost studied in TCR-mediated exhaustion of mouse CD8 T cells in context of tumor or chronic infection^6, 13, 14^. A recent study reported TOX expression in functional circulating human CD8 T_mem_, suggesting TOX expression does not necessarily dictate dysfunction^17, 18^, which led to the speculation that TOX may have distinct roles across species, specifically mice and humans^19^. Alternatively, TOX expression heterogeneity in humans may simply reflect the more complex environment that human T cells are exposed to in every day life, that may not be readily appreciable in specific pathogen-free mice, such as routine inflammatory events in barrier tissues. Thus, we asked if pro-inflammatory cues could be sufficient to increase TOX expression and contribute to TOX heterogeneity. While inflammation has been previously shown to enhance TCR-mediated TOX upregulation (in a VEGF-A-dependent manner that necessitates initial TCR signaling)^48^, our findings are, to the best of our knowledge, the first to demonstrate TOX expression in the absence of agonist TCR signals. Transient IL-12/15/18 and TCR stimulation increased PD-1 and TOX expression in most CD8 T_mem_. In mouse, dysfunctional P14 T_mem_ from LCMV Docile infected mice still increased surface PD-1 expression after TCR stimulation, while IL-12/15/18 had little to no effect on TOX expression. Similarly, human T_EMRA_ showed limited to no increase in TOX expression following exposure to IL-12/15/18. The underlying mechanisms will require further investigation, but one could speculate that the cytokine stimulation is simply not potent enough to further enhance the already ongoing effector or activation program in these two memory T cell subsets. The notion that TOX, but also PD-1 expression can indicate an ongoing effector or activation program in CD8 T cells is important, since PD-1 and (now also) TOX are used as biomarkers of T cell exhaustion^49, 50, 51^. Of note, certain features of general activation programs of CD8 T_mem_ appear to be well conserved and have also been reported as transcriptomic overlap of tissue-resident, recently-activated, and exhausted CD8 T cells^52^. While infection parameters and inflammatory events are well defined in mouse model studies, most human studies remain agnostic in regard to the infection and activation history of Ag-specific T cells. This in turn makes it difficult to correctly interpret the underlying reason for expression of PD-1 and TOX by human T cells.

Our data emphasize the need for conservative interpretation of TOX in regard to activation and exhaustion and also caution against interpreting TOX expression purely through the lens of recent TCR-mediated activation. TOX expression has been predictive of T cell exhaustion and unfavorable outcome in hepatocellular carcinoma animal models and clinical samples^53^, in line with the paradigm of TOX-mediated TCR-dependent T cell dysfunction. However, other studies have yielded contradictory data. Meta analyses of TOX expression in breast cancers reported TOX levels paradoxically correlating with increased immune cell function and favorable prognosis^54^. This is perplexing, as in tumors, TOX expression is associated with T cell dysfunction^6, 13, 14^. This discrepancy could in part be explained by TOX upregulation during activation, akin to what we observed during T cell activation in TCR-dependent and -independent stimulations. Thus, our data stress that all possible activation pathways of TOX and PD-1 induction must be considered before interpreting TOX as a biomarker of T cell dysfunction. A well done human study that interrogated TOX heterogeneity found elevated TOX in CMV- and EBV-specific CD8 T_mem_ and hypothesized recent viral reactivation provided cognate Ag to facilitate TCR-mediated upregulation of TOX. This is certainly a plausible explanation, but our data highlight the need to also consider recent exposure to inflammation as a critical parameter affecting TOX expression. Conventional CD8 T_EM_ and T_EMRA_ (the predominant phenotype of CMV- and EBV-specific CD8 T cells) express elevated levels of TOX basally, T_CM_ (including IAV-specific CD8 T cells) and innate-like MAIT cells can, too, upregulate TOX expression following inflammation-mediated activation. Importantly, our data highlight that this mechanism of TOX expression is conserved across species, conventional CD8 T_mem_ subsets, and innate-like MAIT cells.

Since pro-inflammatory cytokines can concurrently induce TOX and PD-1 expression, these signals may drive TOX heterogeneity in other contexts. P14 tissue-resident memory T cells showed increased *Tox* expression at homeostasis, which has been observed 90 days post priming with LCMV Armstrong^20^. Since the acute infection is cleared well before this timepoint, it is unlikely that continued TCR signaling by cognate Ag drives this phenotype, despite elevated transcripts encoding mediators of TCR signaling^20^. IL-15, however, is likely present within the tissue microenvironment. IL-15 has been implicated in T_RM_ maintenance^25, 26^, and transcriptional profiles indicative of IL-15/STAT5 signaling can be detected in human T_RM_^23, 24^. Thus, IL-15 in tissue microenvironments may also contribute to TOX heterogeneity. Future work will be necessary to dissect the role of these inflammatory cues versus other signals that can shape T_RM_ phenotype, such as co-stimulation and tonic TCR signaling^21^.

Previous studies have demonstrated that TOX ablation or knockdown leads to PD-1 downregulation in models of exhaustion^6, 13, 15, 47^, and conversely, introduction of TOX-expressing constructs enhances PD-1 expression^15, 47^. Similarly, our data showed a close correlation in regards to TOX and PD-1 expression levels, but we found that PD-1 expression could be induced in stimulated *Tox^-/-^* P14 T_mem_. Of note, these *Tox^-/-^* P14 T_mem_ lack exon 5, which abrogates the ability to function as a transcription factor, but the truncated protein is still expressed and detected by the TOX antibody. Alfei et al. previously showed that the early wave of effector cells formed from *Tox^-/-^* T_naïve_ expressed significant levels of PD-1 independently of functional TOX. However, TOX was required for the expression of high levels of PD-1 at later stages, once the initial population of exhausted effector T cells had been replaced by a proliferation competent TCF1 progenitor population^31^. Together, these data suggest that long-term expression of PD-1 requires TOX, but activation-induced expression of PD-1 is TOX-independent. In the absence of TOX, PD-1 expression could be driven TOX2, which can induce PD-1 expression in CD8 T cells^15, 47^; however, it remains unclear if TOX2 is also upregulated by transient TCR- or cytokine-mediated stimulation. Similarly, how different activating signals integrate to regulate TOX expression also requires further studies: while inflammatory cues increase TOX expression in memory T cells, increased IL-12 signaling during the priming of T_naïve_ has been shown to limit subsequent TOX expression at steady state^55, 56^.

Overall, our data suggest that the mechanisms that regulate TOX expression, both at homeostasis and after transient TCR or cytokine stimulation, are remarkably similar and quite possibly highly conserved between humans and mice. Our data further highlight the need to consider TOX and PD-1 expression as prominent indicators of ongoing activation and effector programs in T_mem_ instead of exclusive biomarkers of exhaustion.

## Materials and methods

### Mice

Mouse protocols and experimentation conducted at the Fred Hutchinson Cancer Research Center were approved by and in compliance with the ethical regulations of the Fred Hutchinson Cancer Research Center’s Institutional Animal Care and Use Committee. Experiments performed at the Technical University of Munich were in compliance with institutional and governmental regulations in Germany and approved by the veterinarian authorities of the Regierung von Oberbayern in Germany. All animals were maintained in specific pathogen-free facilities and infected in modified pathogen-free facilities. Experimental groups were non-blinded, animals were randomly assigned to experimental groups, and no specific method was used to calculate sample sizes.

We purchased 6-week-old female C67BL/6J mice from the Jackson Laboratory; *Tox^-/-^* P14 mice (P14 *Tox^tm1c(KOMP)Wtsi^;Mx^Cre^;Rosa26-STOP-eYFP)* were generated as previously described^6^. Both WT and *Tox^-/-^* P14 mice, OT-I mice, and gBT-I mice were maintained on CD45.1 congenic backgrounds. We euthanized mice in accordance with institutional protocols and subsequently collected spleens and lymph nodes (LNs) for experimentation.

### Development of memory mice

We prepared a single-cell suspension of LN cells that were harvested from female OT-I, P14, or gBT-I mice by mechanically passing LN tissue through a 70-100μm strainer. To enrich transgenic T cells, we used MACS with a CD8 negative selection kit (Miltenyi Biotec).

For OT-I memory mice, we adoptively transferred 1□×□10^4^ OT-I T cells in sterile 1× PBS i.v. per C57BL/6J recipient, and subsequently infected recipients i.v. with 1-2 × 10^7^ PFU OVA-expressing vesicular stomatitis virus (VSV-OVA) or 4 × 10^3^ CFU OVA-expressing *Listeria monocytogenes* (LM-OVA). For gBT-I memory mice, we adoptively transferred 5 × 10^4^ gBT-I T cells i.v. and subsequently infected recipient mice i.v. with or 4 × 10^3^ CFU herpes simplex virus 2 (HSV2) glycoprotein B (gB)-expressing *L. monocytogenes* (LM-gB). We allowed ≥ 60 days to pass after initial VSV or LM infections before assaying tissues.

For P14 memory mice, we adoptively transferred 2 × 10^3^ WT P14 T cells i.v. and subsequently infected recipient mice i.v. with 2 × 10^5^ PFU LCMV Armstrong clone (LCMV Arm.) or 2 × 10^6^ PFU LCMV Docile clone (LCMV Doc.). For *Tox^-/-^* P14 memory mice, we adoptively transferred 2 × 10^3^ *Tox^-/-^* P14 memory mice and subsequently infected with 2 × 10^5^ PFU LCMV Arm.; we allowed 28 days to pass after initial LCMV infection before assaying tissues.

### Human PBMC and study approval

Twenty-three healthy, HIV-uninfected adults were recruited by the Seattle HIV Vaccine Trials Unit (Seattle, Washington, USA) as part of the study “Establishing Immunologic Assays for Determining HIV-1 Prevention and Control.” These samples are also known as the Seattle Area Control (SAC) Cohort. All participants were provided and signed informed consent, and the Fred Hutchinson Cancer Research Center Institutional Review Board approved the study protocol.

### T cell isolation and in vitro stimulation

We harvested spleen and LN from memory mice and mechanically prepared single-cell suspensions. We thawed ~4 × 10^7^ cryopreserved PBMC in human RP10 media (RPMI1640 supplemented with 10% FBS, 2mM L-glutamine, 100 U/mL penicillin-streptomycin). To enrich bulk T cells from single cell suspensions, we respectively used mouse- and human-specific T cell negative isolation MACS (STEMCELL Technologies, Canada). We plated 0.5–1 × 10^6^ T cells per well in 96-well V-bottom tissue culture plates. We cultured cells in human RP10 or mouse RP10 media (RPMI 1640 supplemented with 10% FBS, 2mM L-glutamine, 100 U/mL penicillin-streptomycin, 1mM sodium pyruvate, 0.05mM β-mercaptoethanol, and 1mM HEPES). To stimulate cells, we cultured mouse T cells in mouse RP10 with rIL-12, rIL-15, and rIL-18 (BioLegend) (each at 100ng/mL), with Dynabeads mouse T-Activator (Thermo Fisher) anti-CD3/CD28 beads (at a 1:1 bead:cell ratio), or with media alone. For human T cell stimulations, we used human RP10 media with combinations of rIL-6 (BioLegend), rIL-12, rIL-15, and/or rIL-18 (Peprotech) (each at 100ng/mL), with Dynabeads human T-Activator (Thermo Fisher) anti-CD3/CD28 beads (at a 1:1 bead:cell ratio), or with RP10 alone. We cultured cells at 37°C, 5% CO_2_, sampling cells at 0, 24, and 48 hours for flow staining. For intracellular cytokine staining (ICS), we added GolgiPlug (BD Biosciences) at a 1:1,000 dilution 8 hours prior to cell harvest.

### Flow cytometric analysis

We conducted all flow staining for mouse and human T cells on ice and at room temperature, respectively. All mouse and human flow panel reagent information, stain conditions, and gating are included in **(Supplemental Fig. 8-11, Supplemental tables 1-6)**. We conducted LIVE/DEAD fixable aqua or blue viability dye (AViD or BViD, repectively) or Zombie Near-IR viability dye (NIRViD) staining in 1× PBS. For surface staining, we utilized FACSWash (1□× PBS supplemented with 2% FBS and 0.2% sodium azide) as the stain diluent. For all TOX staining panels, we fixed cells with the FOXP3 fixation/permeabilization buffer kit (Thermo Fisher) and conducted intranuclear stains using the FOXP3 permeabilization buffer (Thermo Fisher) as diluent. To minimize day-to-day variation for TOX staining, we conducted all intracellular stains within a batch (0, 24, and 48-hour samples) at the same time. We resuspended cells in FACSWash and acquired events on a FACSSymphony, which we analyzed using FlowJo v10 (BD Biosciences). We conducted statistical testing using Prism v8 (GraphPad).

## Acknowledgements

We thank Andrea Schietinger for helpful discussions and critical review of the manuscript. We also thank the Prlic lab, especially Jami Erickson, Florian Mair, Marie Frutoso, and Veronica Davé for critical review of the manuscript. This work was supported by National Institutes of Health grant R01 AI123323 (to M.P.), National Cancer Institute Grant F99 CA245735 (to N.J.M). N.J.M. is a Leslie and Pete Higgins Achievement Rewards for College Scientists Fellow and Dr. Nancy Herrigel-Babienko Memorial Scholar. D.Z. and J.B were supported by a European Research Council consolidator grant (ToCCaTa) and by the German Research Foundation (SFB1054 and SFB1371).

