## Supplemental Figure 1 for "Inflammatory signals are sufficient to elicit TOX expression in mouse and human CD8 T cells"

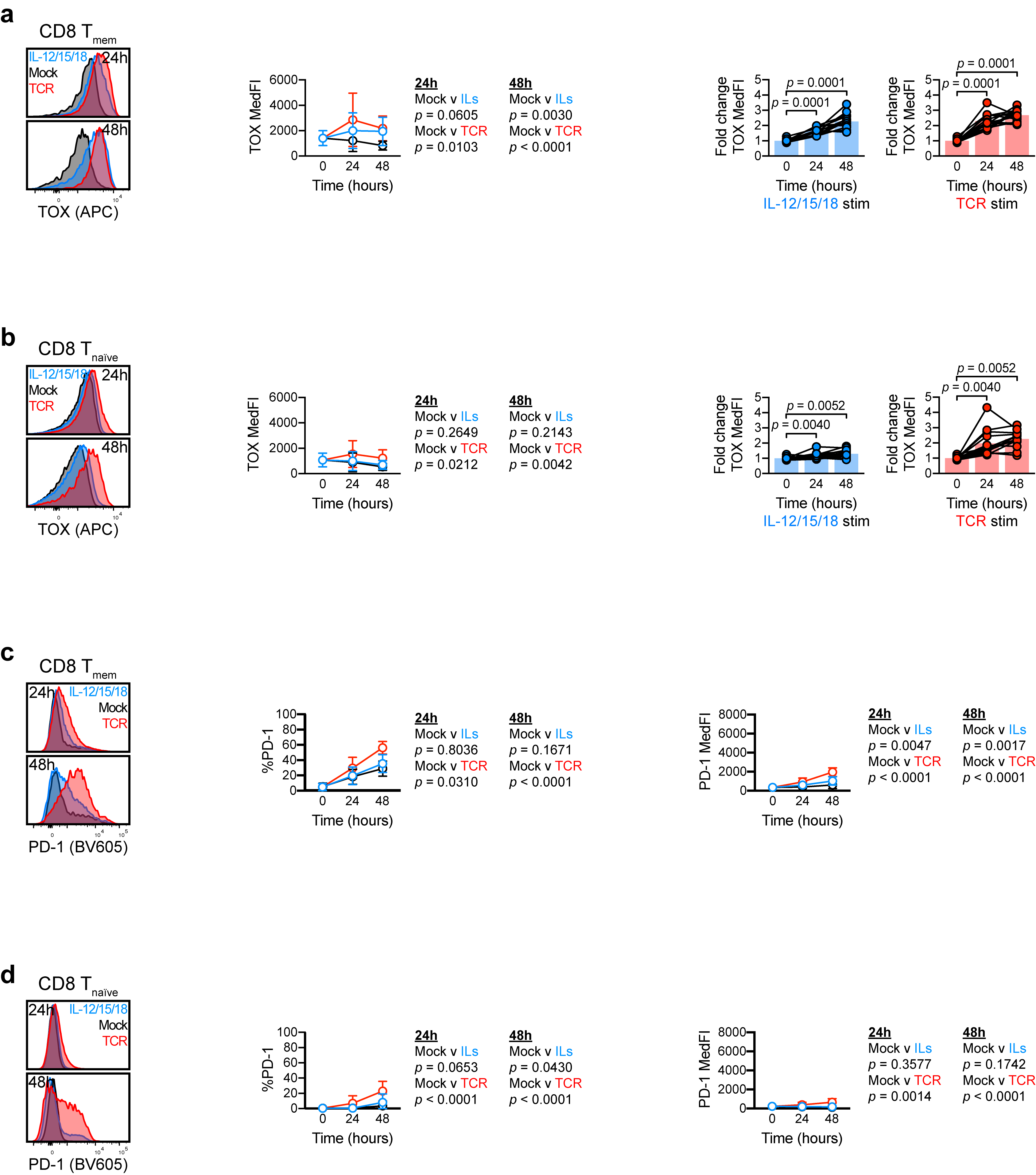

**Supplemental Fig. 1, related to Fig. 1: Cytokine stimulation induces TOX expression in murine CD8 T_mem_**

**a**-**d** T cell stimulation responses after culture with media (mock, black), IL-12/15/18 (each at 100ng/mL, blue), or TCR agonist (1:1 bead to cell ratio, red).  **a**-**b** Stimulation-induced changes of TOX expression in **a** endogenous CD8 T_mem_ and **b** endogenous CD8 T_naïve_ sourced from VSV-OVA OT-I memory mice. **c**-**d** Stimulation-induced changes of PD-1 expression in **c** endogenous CD8 T_mem_ and **d** endogenous CD8 T_naïve_ sourced from VSV-OVA OT-I memory mice. TOX MedFI fold changes in **a** and **b** were calculated against average TOX MedFI from mock stimulations in a subset-specific, batch-specific, and timepoint-specific manner. In **a**-**d**, symbols in line plots comparing stimulation conditions represent the mean across all animals for a specific timepoint/condition ± SD; the indicated statistical significances were calculated using Mann-Whitney tests. In **a** and **b**, bar chart symbols represent one animal at a unique timepoint/condition and are connected by animal identity, with bar indicating mean; the indicated statistical significances were calculated using Wilcoxon matched-pairs signed rank tests. All figures depict results from *n* = 14 mice across 7 experiments. All representative flow plots are sourced from the same animal.
