## Supplemental Figure 2 for "Inflammatory signals are sufficient to elicit TOX expression in mouse and human CD8 T cells"

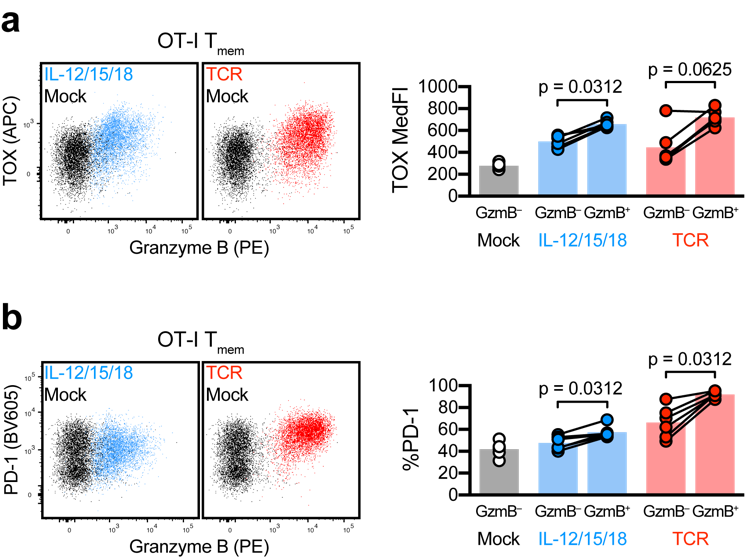

**Supplemental Fig. 2, related to Fig. 2: TOX and PD-1 expression occur in functional CD8 T cells**

T cells were isolated from VSV-OVA OT-I ­memory mice and stimulated (mock, black; IL-12/15/18, blue; TCR, red) for 24 hours (schematic shown in **Fig. 2a**). **a-b** Expression of **a** TOX and **b** PD-1 within granzyme B (GzmB)-positive and -negative OT-I T_mem_ after stimulation. Symbols in **a** and **b** represent a T cell population within a unique animal with symbols connected by animal identity (*n* = 6 across 2 experiments). Bars represent mean and indicated statistical significances were calculated by Wilcoxon matched-pairs signed rank test.
