## Supplemental Figure 3 for "Inflammatory signals are sufficient to elicit TOX expression in mouse and human CD8 T cells"

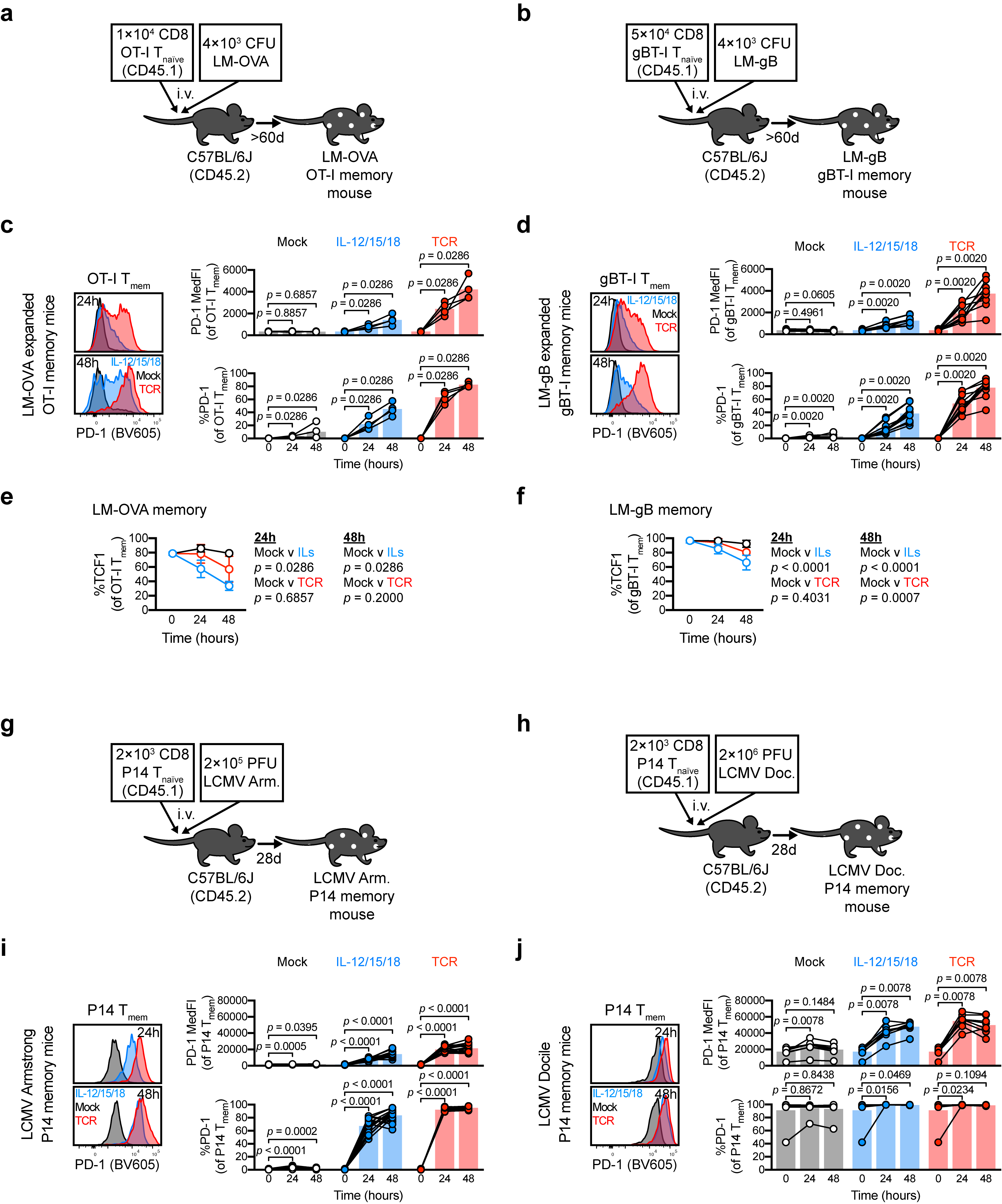

**Supplemental Figure 3, related to Fig. 3: TOX induction varies across memory T cells elicited by different infections**

**a-f** Changes in PD-1 and TCF1 expression within LM-expanded TCR transgenic T_mem_. **a-b** Experiment schematic outlining the expansion of **a** OT-I (OVA-specific) and **b** gBT-I (gB-specific) transgenic T cells with LM-OVA and LM-gB, respectively. MACS-enriched T cells from these memory mice were stimulated with media alone (mock), recombinant IL-12, -15, and -18 in combination (IL-12/15/18) (each at 100ng/mL), or anti-CD3/CD28 microbeads (TCR) at a ~1:1 cell:bead ratio. **c, d** PD-1 MedFI and expression frequency within stimulated **c** OT-I T_mem_ or **d** gBT-I T_mem_. **e, f** TCF1 expression within stimulated **e** OT-I and **f** gBT-I T_mem_. **g-j** Changes in PD-1 expression within LCMV Armstrong- and Docile-expanded P14 T_mem_. **g-h** Experiment schematic outlining the expansion of P14 transgenic T cells with **g** LCMV Armstrong or **h** LCMV Docile, which respectively cause acute and chronic infection. **i-j** PD-1 MedFI and expression frequency within **i** LCMV Armstrong- and **j** Docile-expanded P14 T_mem_. Symbols in **c, d, i, j** represent T cell populations from a single animal at a unique timepoint/condition and are connected by matched donor identities (when applicable), with bars depicting mean. Symbols in **e, f** represent mean values ± SD. Indicated statistical significances in **c, d, i, j** were calculated by Wilcoxon matched-pairs signed rank tests, and those in **e, f** were calculated using Mann-Whitney tests. Data in **a-f** depict *n* = 4 LM-OVA OT-I memory mice and *n =* 10 LM-gB gBT-I memory mice across 2 experiments. Data in **g** and **j** depict *n* = 17 LCMV Armstrong expanded P14 memory mice across 4 experiments and *n* = 8 LCMV Docile expanded P14 memory mice across 2 experiments.
