## Supplemental Figure 4 for "Inflammatory signals are sufficient to elicit TOX expression in mouse and human CD8 T cells"

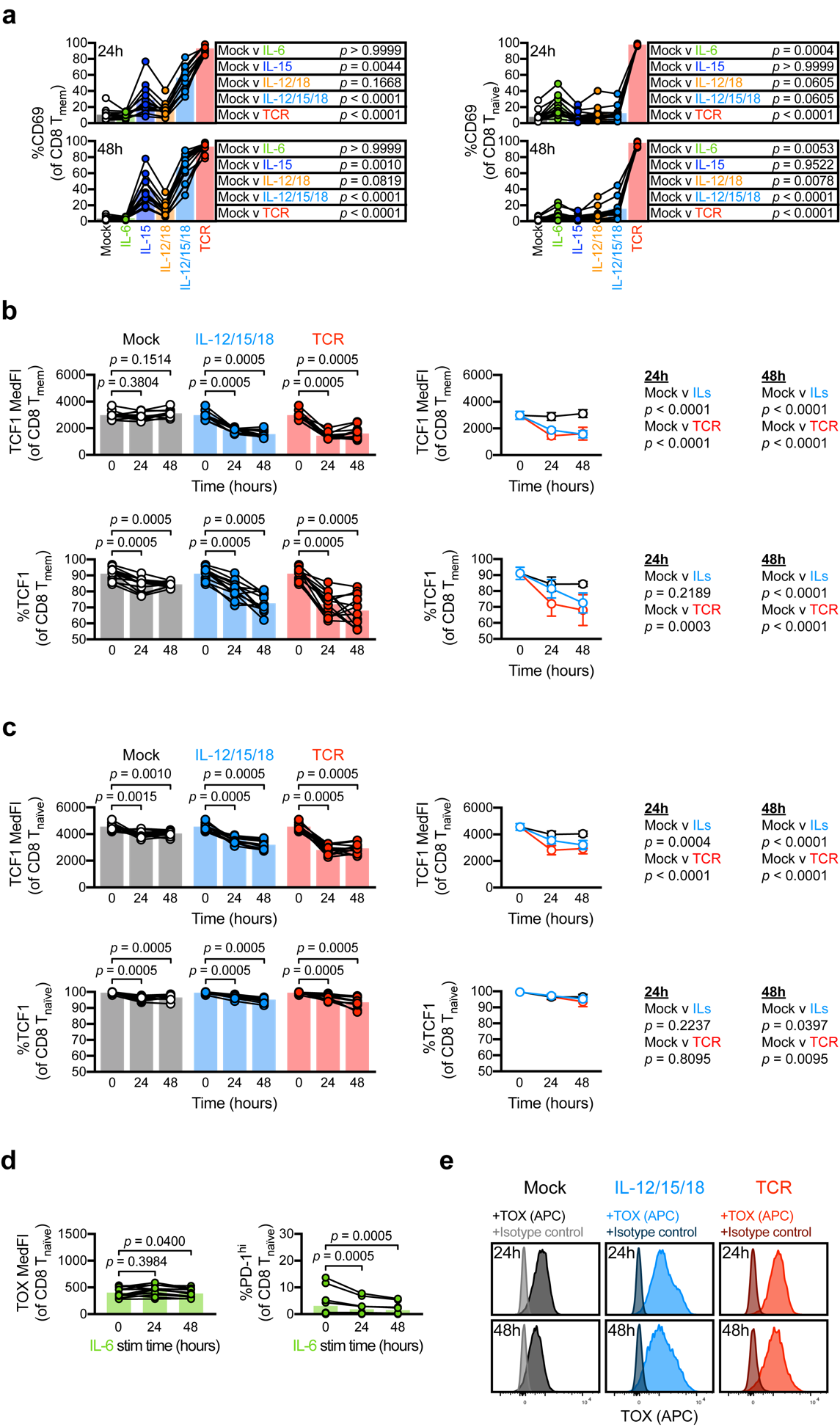

**Supplemental Figure 4., related to Fig. 4: Inflammatory cytokines are potent inducers of TOX and PD-1 in human T_mem_**

**a**-**b** CD69 expression in stimulated CD8 T_mem­_ (left) and CD8 T_naïve_ _­_(right) subsets. **b**-**c** TCF1 MedFI and expression frequency in **b** CD8 T_mem­_ and **c** CD8 T_naïve_ _­_over stimulation time course (mock, black; IL-12/15/18, blue; TCR, red). **d** TOX MedFI and PD-1^hi^ event frequencies in IL-6-stimulated CD8 T_naïve_. **e** TOX and isotype control staining in mock (left), IL-12/15/18 (center), and TCR (right) stimulated CD8 T_mem_. Bar plot symbols in **a**-**d** depict a unique donor at a specific condition/timepoint, with symbols connected by donor identity and bars depicting mean; indicated statistical significances were calculated by **a** Friedman tests with Dunn’s multiple comparisons tests or **b**-**d** Wilcoxon matched-pairs signed rank tests. Line plot symbols in **b** and **c** depict means ± SD across stimulation conditions with indicates statistical significances calculated by Mann-Whitney tests. **a** and **d** represent *n* = 11 donors across 2 experiments; **b** and **c** represent *n* = 12 donors across 2 experiments; **e** represent *n* = 3 donors.
