## Supplemental Figure 5 for "Inflammatory signals are sufficient to elicit TOX expression in mouse and human CD8 T cells"

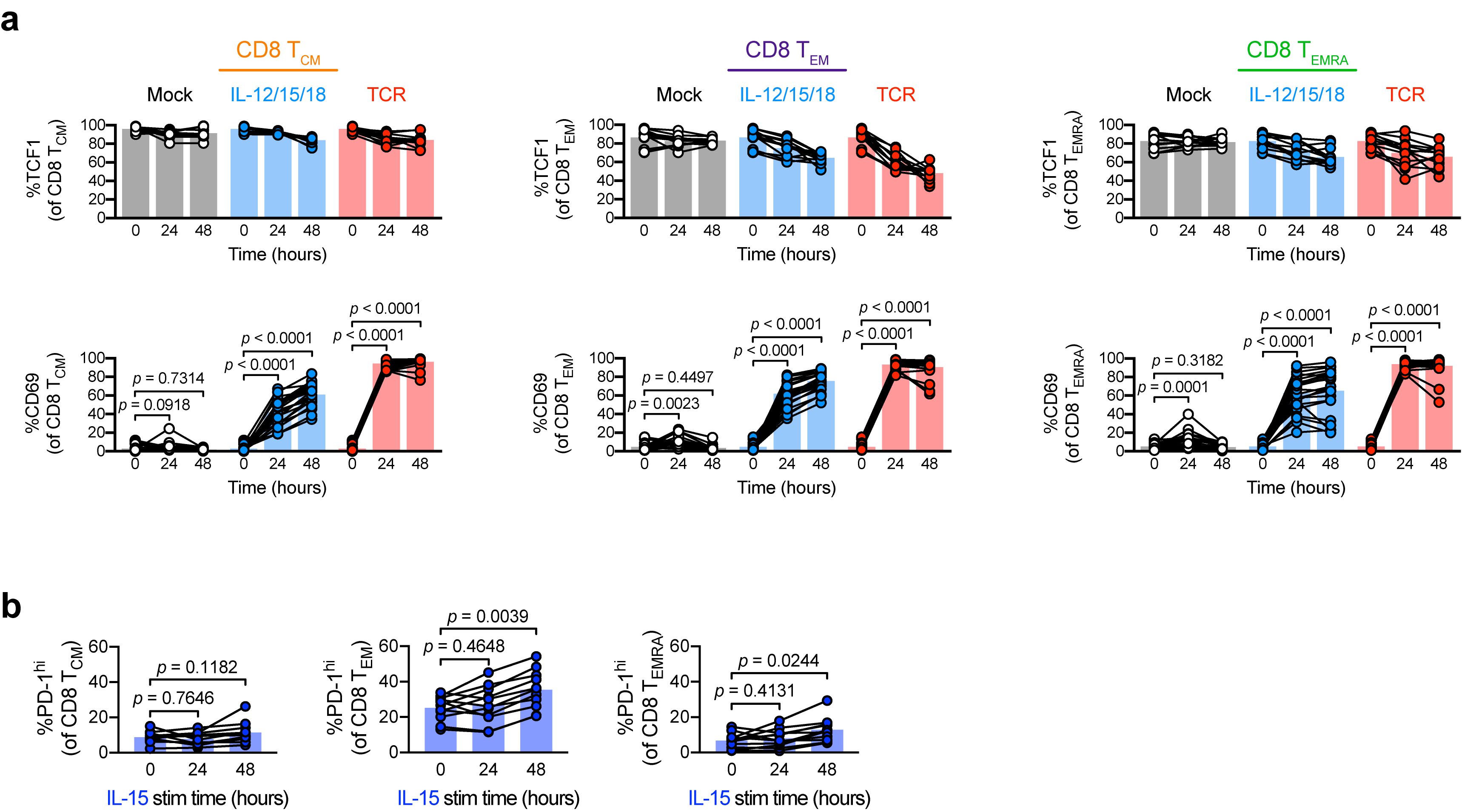

**Supplemental Figure 5, related to Fig. 5: TOX and PD-1 upregulation are largely independent of T_mem_ subset**

**a** TCF1 (top row) and CD69 (bottom row) event frequency within CD8 T_mem_ subsets after mock (black), IL-12/15/18 (blue), or TCR (red) stimulation. **b** PD-1^hi^ event frequency within CD8 T_mem_ subsets after IL-15 stimulation. All cytokines were at 100ng/mL, each in all stimulation conditions. In **a**-**b**, each symbol represents a cell population within a donor at a unique condition/timepoint, with symbols connected by donor identity and bar representing mean. All indicated statistical significances were calculated by Wilcoxon matched-pairs signed rank tests. Data in **a** depicts 12 (%TCF1) donors over 2 experiments and 23 donors (%CD69) over 4 experiments; **b** depicts 11 donors over 2 experiments.
