## Supplemental Figure 6 for "Inflammatory signals are sufficient to elicit TOX expression in mouse and human CD8 T cells"

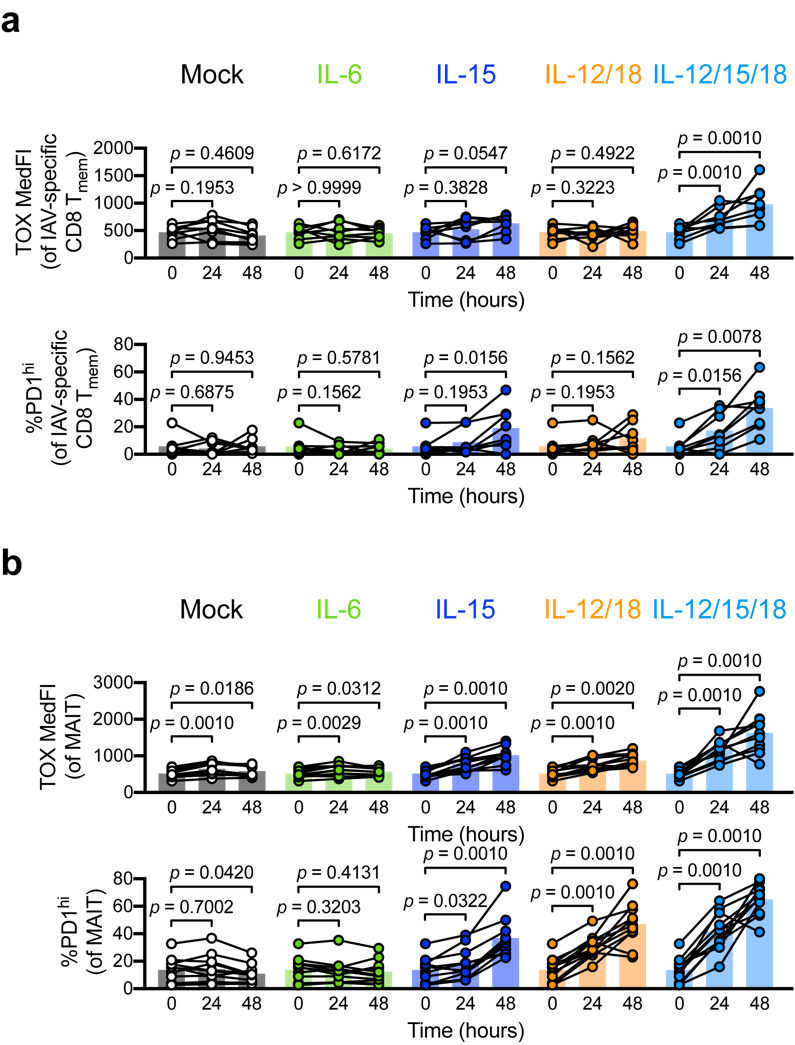

**Supplemental Figure 6, related to Fig. 6: Stimulation induces TOX and PD-1 expression in conventional and innate-like T cells**

**a**-**b** TOX MedFI and frequency of PD-1^hi^ events in **c** IAV-specific CD8 T cells and **d** MAIT cells after mock (black), IL-6 (green), IL-15 (dark blue), IL-12/18 (orange), or IL-12/15/18 (blue) stimulation (all cytokine concentrations 100/ng/mL, each). Data in **a** depicts 8 donors over 2 experiments; **b** depicts 11 donors over 2 experiments. All indicated statistical significances were calculated using Wilcoxon matched-pairs signed rank tests.
