## Supplemental Figure 7 for "Inflammatory signals are sufficient to elicit TOX expression in mouse and human CD8 T cells"

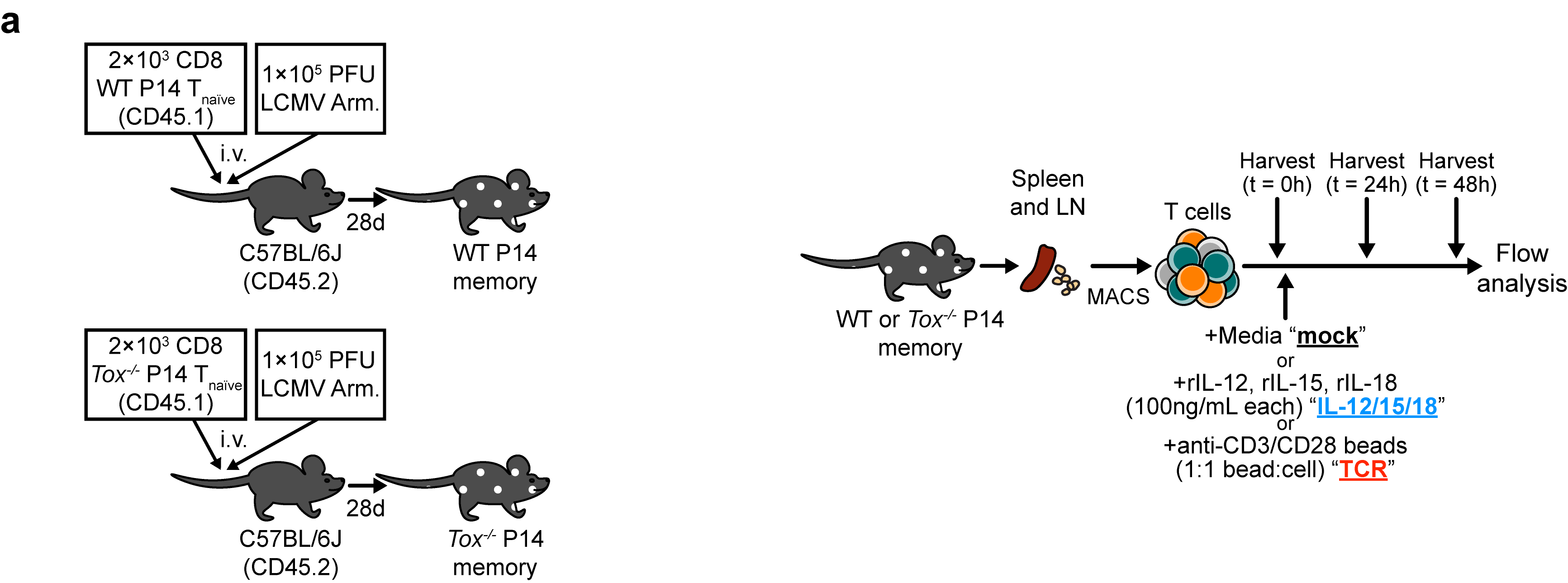
**Supplemental Figure 7, related to Fig. 7: TOX deficiency does not abrogate stimulation-induced PD-1 expression**

**a** Experiment schematic for adoptive transfers, tissue harvests, and stimulations conducted in Fig. 7.
