## Supplemental Figure 8 for "Inflammatory signals are sufficient to elicit TOX expression in mouse and human CD8 T cells"

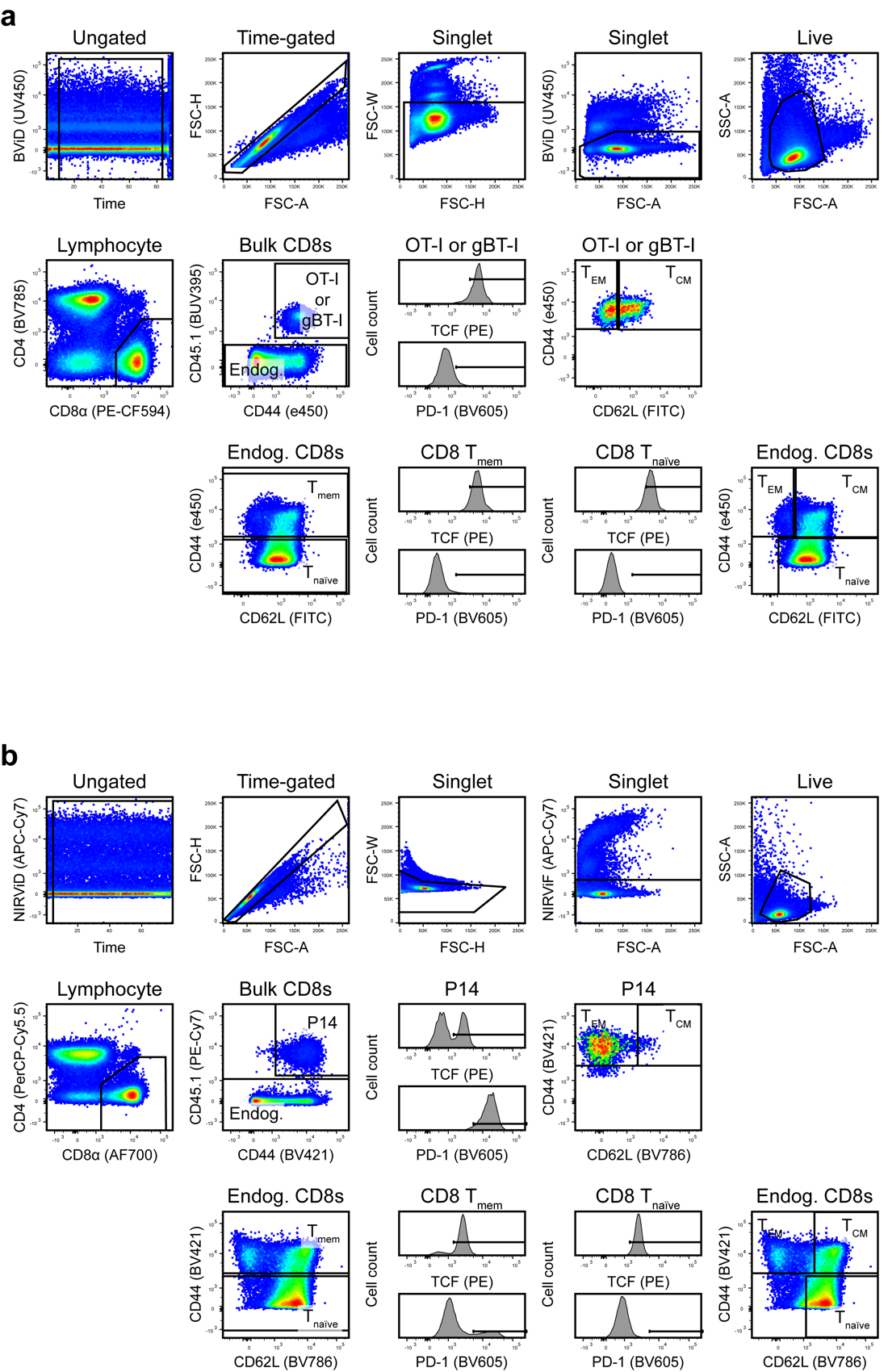

**Supplemental Figure 8, related to Fig. 1, 3, 7, Supplemental Fig. 1, 3**

**a**-**b** Representative flow gating for **a** VSV-OVA OT-I memory mice, LM-OVA OT-I memory mice, and LM-gB gBT-I memory mice (unstimulated, 0 h timepoint, VSV-OVA OT-I memory mouse) and **b** LCMV Armstrong- and Docile-expanded WT and Tox^–/–^ P14 memory mice (unstimulated, 0 h timepoint, LCMV Docile WT P14 memory mouse).
