## Supplemental Figure 9 for "Inflammatory signals are sufficient to elicit TOX expression in mouse and human CD8 T cells"

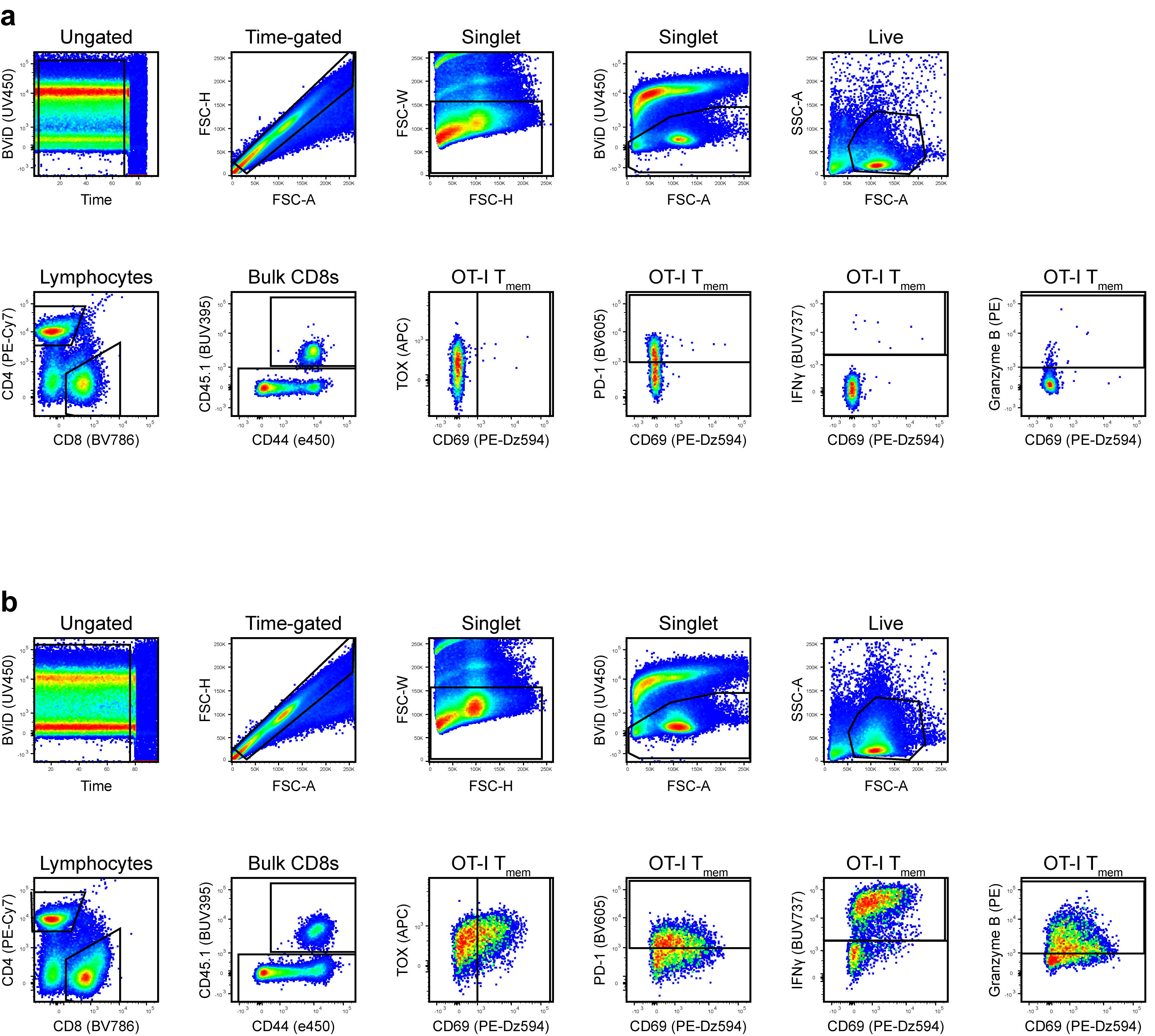

**Supplemental Fig. 9. related to Fig. 2, Supplemental Fig. 2**

**a** and **b** Representative flow gating for intracellular cytokine staining in parallel with TOX and PD-1 interrogation in mouse T cells (**a** mock-stimulated, 24h; **b** IL-12/15/18-stimulated, 24h).
