## Supplemental Figure 10 for "Inflammatory signals are sufficient to elicit TOX expression in mouse and human CD8 T cells"

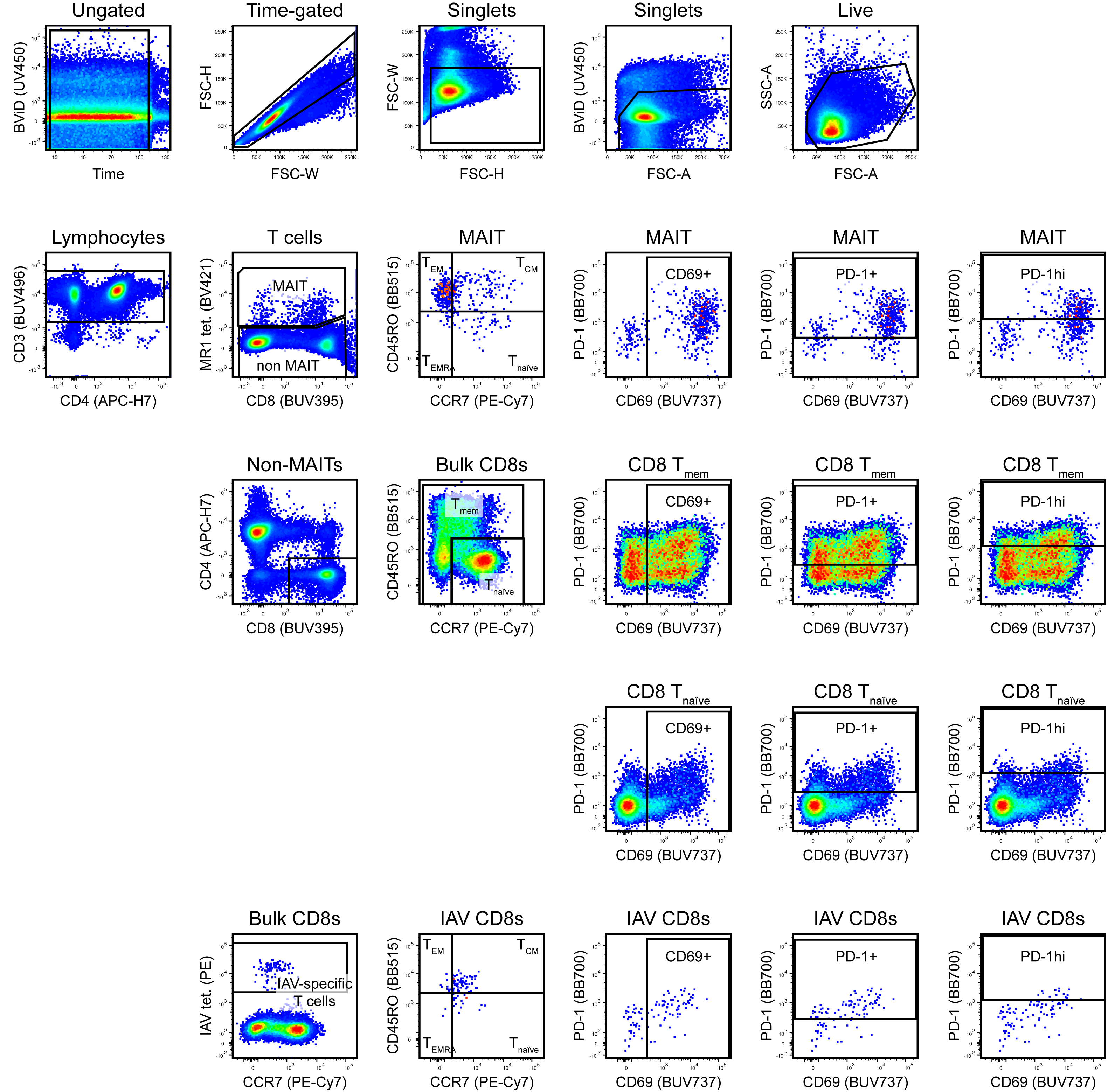

**Supplemental Fig. 10. related to Fig. 4, 5, 6, Supplemental Fig. 4, 5, 6**

**a** Representative flow gating for TOX and PD-1 interrogation in human T cells with IAV and MR1 tetramers (IL-12/15/18-stimulated, 24h).
