## Supplemental Figure 11 for "Inflammatory signals are sufficient to elicit TOX expression in mouse and human CD8 T cells"

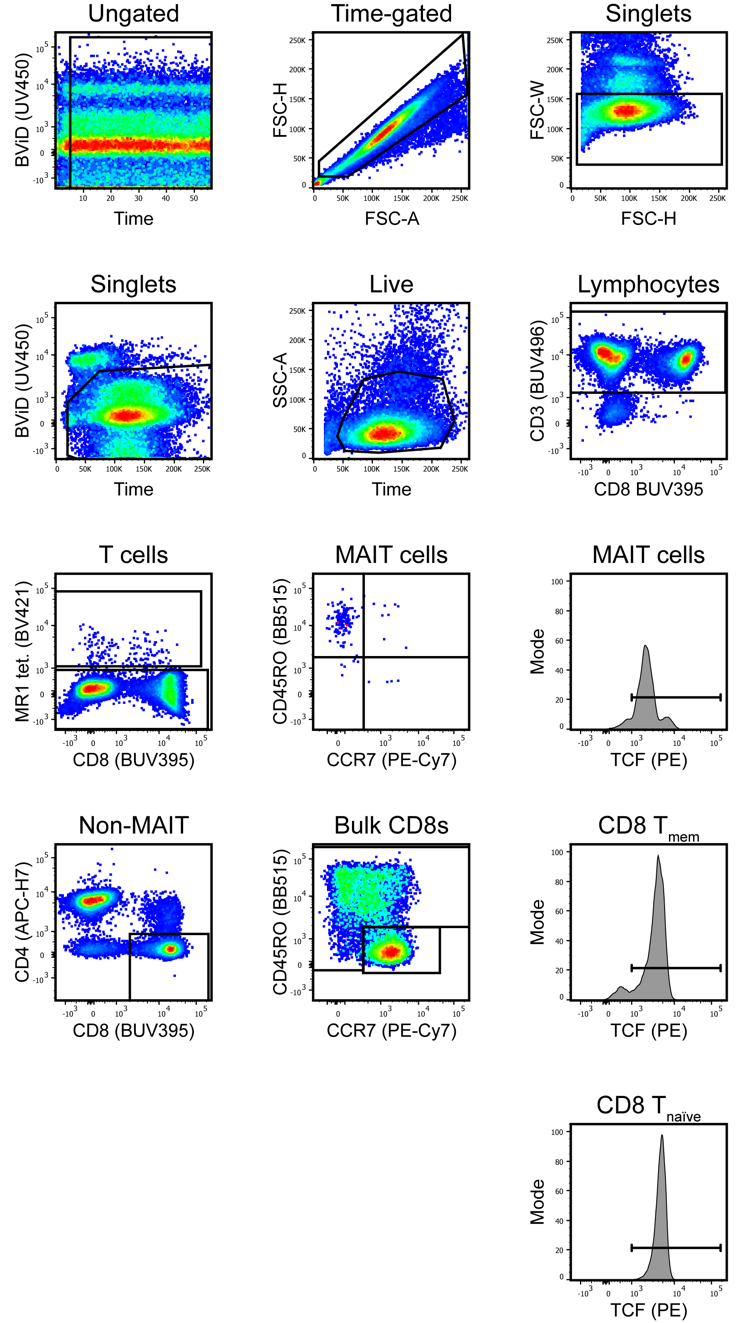

**Supplemental Fig. 11. related to Fig. 5, Supplemental Fig. 4, 5**

**a** Representative flow gating for TOX, PD-1, and TCF1 interrogation in human T cells (unstimulated, 0h).
