## Supplemental Table 1 for "Inflammatory signals are sufficient to elicit TOX expression in mouse and human CD8 T cells"

| Reagent | Fluor | Clone | Vendor | Dilution |
| --- | --- | --- | --- | --- |
| Viability stain (1x PBS diluent, 20 min, on ice) | | | | |
| Zombie NIR fixable viability kit  (NIRViD) | APC-Cy7 | NA | BioLegend | 1:500 |
| Surface stain (FACSWash diluent, 30 min, on ice) | | | | |
| Fc block  (CD16/CD32) | Unconjugated | 2.4G2 | BD | 1:200 |
| CD4 | PerCP-Cy5.5 | RM4-4 | BioLegend | 1:200 |
| CD8α | AF700 | 53-6.7 | Thermo Fisher | 1:200 |
| CD45.1 | PE-Cy7 | A20 | Thermo Fisher | 1:200 |
| CD44 | BV421 | IM7 | BioLegend | 1:200 |
| PD-1 | BV605 | 29F.1A12 | BioLegend | 1:100 |
| CD62L | BV785 | MEL-14 | BioLegend | 1:200 |
| Fix (1x eBioscience FOXP3 fixation buffer, 20 min, on ice) | | | | |
| Intracellular stain (1x eBioscience FOP3 permeabilizaiton buffer diluent, 30 min, on ice) | | | | |
| Fc block  (CD16/32) | Unconjugated | 2.4G2 | BD | 1:200 |
| TOX | e660 | TXRX10 | Thermo Fisher | 1:200 |
| TCF1/7 | PE | S33966 | BD | 1:200 |

**Supplemental Table 1.** Mouse flow cytometry panel for SPF C57BL/6J, LCMV Armstrong P14 memory mice, and LCMV Docile P14 memory mice.
