## Supplemental Table 2 for "Inflammatory signals are sufficient to elicit TOX expression in mouse and human CD8 T cells"

| Reagent | Fluor | Clone | Vendor | Dilution |
| --- | --- | --- | --- | --- |
| Viability stain (1x PBS diluent, 20 min, on ice) | | | | |
| LIVE/DEAD fixable blue viability dye  (BViD) | UV450 | NA | Thermo Fisher | 1:500 |
| Surface stain (FACSWash diluent, 30 min, on ice) | | | | |
| Fc block  (CD16/CD32) | Unconjugated | 2.4G2 | BD | 1:200 |
| CD4 | BV786 | GK1.5 | BD | 1:200 |
| CD8α | PE-CF594 | 53-6.7 | BD | 1:200 |
| CD45.1 | BUV395 | A20 | BD | 1:200 |
| CD44 | e450 | IM7 | Thermo Fisher | 1:200 |
| PD-1 | BV605 | 29F.1A12 | BioLegend | 1:100 |
| CD62L | FITC | MEL-14 | Thermo Fisher | 1:200 |
| Fix (1x eBioscience FOXP3 fixation buffer, 20 min, on ice) | | | | |
| Intracellular stain (1x eBioscience FOP3 permeabilizaiton buffer diluent, 30 min, on ice) | | | | |
| Fc block  (CD16/32) | Unconjugated | 2.4G2 | BD | 1:200 |
| TOX | APC | REA473 | Miltenyi Biotec | 1:100 |
| TCF1/7 | PE | C63D9 | Cell Signaling | 1:40 |

**Supplemental table 2.**  Mouse flow cytometry panel for VSV-OVA OT-I memory mice, LM-OVA OT-I memory mice, and LM-gB gBT-I memory mice.
