## Supplemental Table 5 for "Inflammatory signals are sufficient to elicit TOX expression in mouse and human CD8 T cells"

| Reagent | Fluor | Clone | Vendor | Dilution |
| --- | --- | --- | --- | --- |
| Viability stain (1x PBS diluent, 20 min, room temperature) | | | | |
| LIVE/DEAD fixable blue viability dye  (BViD) | UV450 | NA | Thermo Fisher | 1:500 |
| Tetramer stain (FACSWash diluent, 60 min, room temperature) | | | | |
| MR1 OP-5-RU (MAIT cell) tetramer | BV421 | NA | NIH Tetramer Core | 1:500 |
| HLA-A*02 influenza A virus (IAV) (GILGFVFTL) tetramer | PE | NA | Fred Hutch Immune Monitoring | 1:150 |
| TruStain FcX  (Fc block) | NA | NA | BioLegend | 1:20 |
| Surface stain (FACSWash diluent, 20 min, room temperature) | | | | |
| CD3 | BUV496 | UCHT1 | BD | 1:40 |
| CD4 | APC-H7 | RPA-T4 | BD | 1:40 |
| CD8 | BUV395 | RPA-T8 | BD | 1:80 |
| CD45RO | BB515 | UCHL1 | BD | 1:160 |
| CD69 | BUV737 | FN50 | BD | 1:40 |
| CCR7 | PE-Cy7 | 3D12 | BD | 1:40 |
| PD-1 | BB700 | EH12.1 | BD | 1:20 |
| Fix (1x eBioscience FOXP3 fixation buffer, 20 min, room temperature) | | | | |
| Intracellular stain (1x eBioscience FOP3 permeabilizaiton buffer diluent, 30 min, room temperature) | | | | |
| TOX* | APC | REA473 | Miltenyi Biotec | 1:80 |
| KLH-specific REA control antibody I* | APC | REA293 | Miltenyi Biotec | 1:80 |

**Supplemental table 5.** Flow cytometry panel for HLA-A*02 human PBMC samples.

*Reagents were not used in unison, but as stains in separate samples.
